# Engineering a human-based translational activator for targeted protein expression restoration

**DOI:** 10.1101/2025.07.09.663984

**Authors:** Riley W. Sinnott, Ani Solanki, Anitha P. Govind, William N. Green, Bryan C. Dickinson

## Abstract

Therapeutic modalities to programmably increase protein production are in critical need to address diseases caused by deficient gene expression via haploinsufficiency. Restoring physiological protein levels by increasing translation of their cognate mRNA would be an advantageous approach to correct gene expression, but has not been evaluated in an *in vivo* disease model. Here, we investigated if a translational activator could improve phenotype in a Dravet syndrome mouse model, a severe developmental and epileptic encephalopathy caused by *SCN1a* haploinsufficiency, by increasing translation of the SCN1a mRNA. We identifiy and engineere human proteins capable of increasing mRNA translation using the CRISPR-Cas Inspired RNA-targeting System (CIRTS) platform to enable programmable, guide RNA (gRNA)-directed translational activation with entirely engineered human proteins. We identify a compact (601 amino acid) CIRTS translational activator (CIRTS-4GT3), that can drive targeted, sustained translation increases up to 100% from three endogenous transcripts relevant to epilepsy and neurodevelopmental disorders. AAV-delivery of CIRTS-4GT3 targeting SCN1a mRNA to a Dravet syndrome mouse model led to increased SCN1a translation and improved survivability and seizure threshold - key phenotypic indicators of Dravet syndrome. This work validates a new strategy to address *SCN1a* haploinsufficiency and emphasizes the preclinical potential translational activation has to address neurological haploinsufficiency.

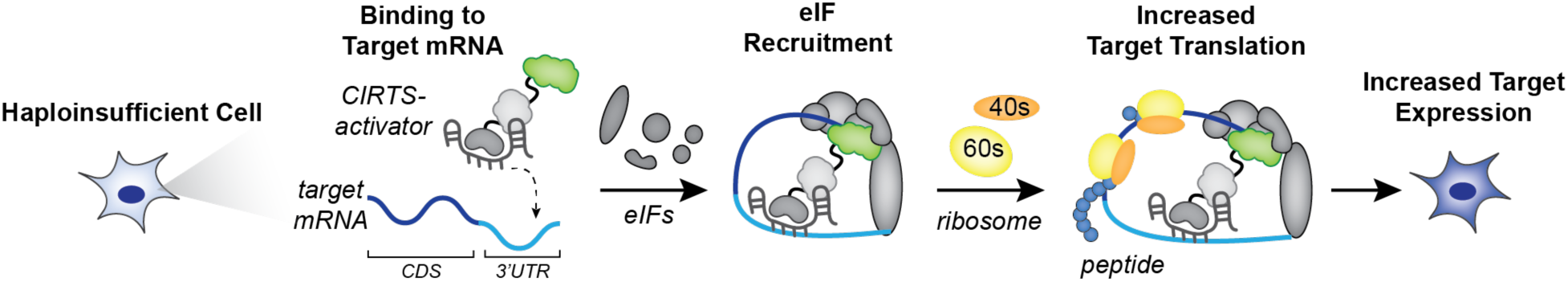

## Introduction

Genetic haploinsufficiency results from a loss-of-function mutation to one gene copy thereby reducing functional protein expression by approximately half (1). Haploinsufficiency to any one of approximately 3,000 identified “dosage” sensitive genes drive diverse diseases, including metabolic disorders, heart disease, cancers and neurological disorders (2,3). Brain-related genes are the most sensitive to aberrant gene dosage (4), and the number of validated causal haploinsufficiency genes identified for epilepsies (5) and neurodevelopmental disorders (6,7) is rapidly growing. While a panoply of existing therapeutic technologies can inhibit (8), knock down (9), knock out (10), and degrade (11) over-active gene products, comparably few are capable of correcting gene under-expression, with even fewer deployable in neurological tissues, presenting a critical unmet therapeutic need for millions of patients (12).

Current therapeutic approaches for restoring homeostatic protein levels from deficient genes typically involve gene supplementation, gene editing, transcriptional activation, suppressor tRNAs, small molecule protein stabilization, or ASOs for splice-switching, lncRNA degradation or disrupting negative regulatory elements (13–16). While each approach is promising in certain contexts, these methods face challenges for targeting haploinsufficiency generally in the brain, such as gene size or oligo-related delivery complications, a significant risk for permanent off-target genomic changes or inducing gene overexpression in off-target cell types eliciting toxicity, or a dependence on target-specific regulatory pathways (**Supplementary Fig. S1**). In contrast, targeting translation by upregulating protein production from endogenous mRNAs using a programmable translational activator offers potential advantages, including no size limitations on the targeted gene, protein activation levels that correspond to the mRNA presence and abundance within each cell, aligning with cell-specific needs, and activation levels that are predictable and governed by the regulatory pathways being utilized (**Supplementary Fig. S1**). Furthermore, the transcriptome-level specificity afforded by RNA-targeting is particularly advantageous in the brain where incredible cell-type diversity complicates DNA-targeting therapeutic delivery (17–19).

While the clinical potential of programmable translational activation for restoring haploinsufficient gene product expression is enticing, especially in the brain, the ability to target translation to relieve haploinsufficient phenotypes remains unproven. We previously established the CRISPR/Cas-inspired RNA-targeting system (CIRTS) - a compact, programmable gRNA-dependent RNA regulatory technology assembled from human proteins to circumvent the immunogenicity and size limitations of CRISPR systems (20–22). Here, we deployed CIRTS to investigate whether activating the translation of a haploinsufficient mRNA was a viable strategy to restore homeostatic gene expression in an animal model of disease (**Fig. 1**). We targeted SCN1a, which is translated into the voltage-gated sodium channel Na_V_1.1, and whose haploinsufficiency results in Dravet syndrome (DS), a severe, early-onset developmental and epileptic encephalopathy (DEE) (23–25). DS is the most common DEE with an incidence of 1:15,500 with SCN1a haploinsufficiency implicated in more than 80% of cases and is resistant to management by conventional anti-epilepsy drugs with no approved cure (26). Beyond the high clinical need for SCN1a-related therapeutics, increasing Na_V_1.1 expression by translational activation could overcome outstanding challenges with known toxicity resulting from *SCN1a* gene therapy in non-target cell types and ongoing work to identify minor cell populations impacted by *SCN1a* haploinsufficiency missed by current delivery methods (27–29). Collectively, this makes DS a powerful test case to develop and assess programmable translational activators.

**Figure 1.**
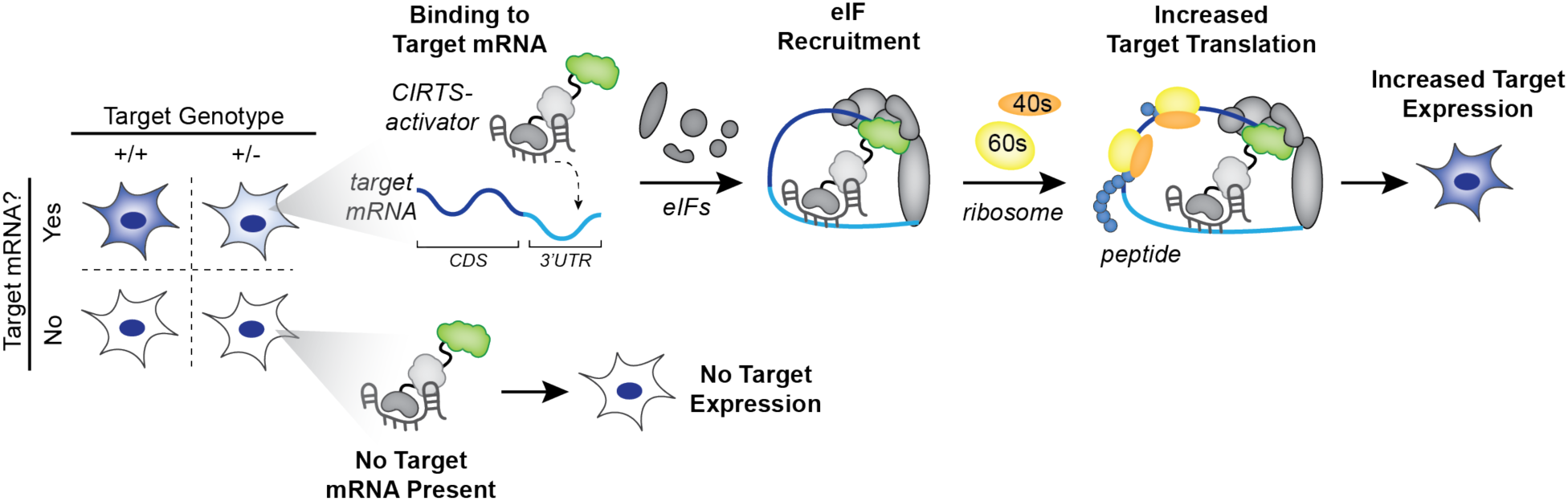
CIRTS-based translational activators (CIRTS-activators) increase target mRNA translation. Representation of CIRTS-activator mediated protein expression increase that occurs only in cells that natively express the target mRNA. CIRTS-activators programmably bind to a target mRNA via the complementary gRNA sequence, then likely recruits eukaryotic initiation factors (eIFs) which in turn recruit the ribosome to promote translation. In the case of cells haploinsufficent for the target, this increase in translation recues target under-expression in a target mRNA expression dependent manner.

In this work, we used CIRTS to screen putative human translational effectors for their ability to activate SCN1a translation, then optimized and minimized a CIRTS translational activator (CIRTSa) construct, CIRTS-4GT3, that increases Na_V_1.1 expression approximately 100% *in cellulo* with a size of only 601 amino acids. We then delivered the optimized SCN1a-targeting CIRTS to *SCN1a* haploinsufficient mice via AAV9, which increased neuronal Na_V_1.1 protein levels and resulted in significantly improved survival and diminished susceptibility to thermal-induced seizures - two hallmarks of the disease pathology. Finally, we also deployed CIRTS-4GT3 to increase expression from two more haploinsufficiency-related mRNA, CHD2 and ARID1B, associated with developmental delay and epilepsy, or intellectual disability and autism spectrum disorder respectively (30–32). Collectively, this work establishes CIRTS-4GT3 as an optimized, programmable translational activator and validates translational activation is a viable strategy to restore gene expression in a neurological haploinsufficiency.

## Materials and Methods

### Ethical Statement

This research complies with all relevant ethical regulations. Animal studies were approved by the Institutional Animal Care and Use Committee (IACUC) at the University of Chicago (Protocol Numbers 72613 and 72016).

### General Cloning

Gibson Assembly (GA) was used to clone all plasmids used for transient transfection in this study. Primers and gBlocks were ordered from IDT. PCR fragments used for Gibson Assembly were amplified with Q5 DNA Polymerase (NEB). After GA, plasmids were transformed into chemically competent DH10β *E. coli* and after a 1 h outgrowth in 2xYT media, were plated on antibiotic agar plates to select single plasmid sequences. Plasmids were sequenced by the University of Chicago Comprehensive Cancer Center DNA Sequencing and Genotyping Facility. All effector sequences (see **Supplementary Table S3**) were amplified from gBlocks for cloning into CIRTS plasmids. CHD2 3’ UTR region was cloned from gBlock and SCN1a 3’ UTR was amplified from reverse transcription reactions completed with total RNA isolated from Neuro 2a (ATCC) cells. All plasmids used in this study are listed in **Supplementary Table S1** with links to fully annotated vectors and are available upon request. Key plasmids will be made available through Addgene.

### Cloning Viral Vectors

Cloning for lentiviral and AAV transgene plasmids was conducted via restriction enzyme (all from NEB) cloning to replace parts in published vectors with the indicated promoters, or CIRTS protein or gRNA expression cassettes. For lentivirus vectors, lentiCRISPR v2 (addgene #52961) was digested with PacI and NheI to clone gRNA cassettes and XbaI and BamHI for CIRTS protein open-reading frames following manufacturer’s protocols. Ligation was carried out with T4 DNA ligase (NEB) at 16 °C for 16 hours. Ligated plasmid was then transformed into chemically competent NEB stable *E. coli* for plating and antibiotic selection at 30 °C. Similarly, for CIRTS-AAV the pX601-SaCas9-gRNA vector (Addgene #61591) was modified to replace the CMV promoter with hSYN by XbaI and AgeI digestion and SaCas9-gRNA was replaced by CIRTS-4GT3-gRNA by AgeI and EcoRI digestion.

### Mammalian Cell Culture

HEK293T (ATCC) and Neuro-2a (ATCC) were maintained using DMEM (L-glutamine, high glucose, sodium pyruvate, phenol red; Corning) supplemented with 10% fetal bovine serum (FBS; Gibco), and 1% penicillin/streptomycin (Gibco). Cells were grown at 37 °C and 5% CO_2_. For transfections, 1% penicillin/streptomycin was omitted. For all experiments, cells had undergone fewer than 18 passages. All cell lines tested negative for mycoplasma contamination.

### Plasmid Transient Transfection

For experiments in HEK293T, cells were plated on a white-walled 96-well plate (Corning) 24 hours before transfection in a final volume of 80 μL per well and grew to 70% confluency by time of transfection. 12 ng reporter plasmid and 250 ng gRNA-CIRTS plasmids were transfected for each condition. Transfections were completed with 0.5 μL lipofectamine 2000 (Invitrogen) per well and Opti-MEM I Reduced Serum Medium (ThermoFisher Scientific) for dilution of the plasmids and lipofectamine following manufacturer’s instructions. Approximately 24 hours after transfection 40 μL of media was removed from each well and replaced with 40 μL fresh DMEM + 10% FBS.

For experiments in Neuro-2a, cells were plated on a 12-well plate (Corning) 24 hours before transfection in a final volume of 1 mL per well and grew to 70% confluency by time of transfection. 1250 ng gRNA-CIRTS plasmid were transfected for each condition using 1.75 μL Lipofectamine 3000 (Invitrogen) and 2 μL P3000 reagent per well with Opti-MEM I Reduced Serum Medium used for dilution according to manufacturer’s instructions. Media was fully exchanged in each well with 1 mL fresh DMEM + 10% FBS approximately 5 hours after transfection.

### gRNA Sequence Design

A guide RNA (gRNA) sequence represents the reverse-complementary RNA sequence for the desired binding site on the target mRNA. In order to generate gRNA sequences with a higher chance of success for specific target engagement we recommend comparing the results for several available gRNA prediction programs listed below as well as filtering out suggested gRNA that 1) contain a run of 5 or more Uracil as this represents a transcription termination sequence for RNA polymerase III encoded on the DNA; 2) contain no significant internal secondary structure precluding sequence availability to pair with the target as predicted by programs such as RNAfold; 3) contain no significant off-target binding sites in the transcriptome as predicted by NCBI Blast analysis.

Three programs were used in combination to generate gRNA sequence candidates. Candidate gRNA sequences were suggested for a given target mRNA by both Cas13design (where the generated 23 nt gRNA were extended to 40 nt on the target by prioritizing the parameters above) and Soligo tool offered in the Sfold package where the oligo length was set to 40 nts. Finally, regions in the target mRNA with existing structures requiring low energy to open (higher probability that the target region will be unpaired) were predicted with RNAfold.

The two gRNA sequence lists rank ordered by Cas13design and Soligo respectively were compared for overlap with priority given to similar sequences overlapping in each list that were predicted to bind in a low energy target region predicted by RNAfold. For sequences of all gRNA used in this study see **Supplementary Table S2.**

RNAfold(33): http://rna.tbi.univie.ac.at/cgi-bin/RNAWebSuite/RNAfold.cgi Cas13design(34): https://cas13design.nygenome.org/ Soligo(35): https://sfold.wadsworth.org/cgi-bin/soligo.pl

### Dual Luciferase Reporter Assay

48 hours after transfection, luciferase activity was assayed according to protocols previously developed and described by Baker et. al. with only minor alterations^1^. 3x firefly assay buffer and 3x nano luciferase assay buffers were prepared fresh before every experiment and protected from light until used. Nano luciferase salts were prepared fresh weekly and Triton lysis buffer was prepared fresh monthly.

Plates were removed from the incubator and left to cool to room temperature for 15 minutes. After plates had cooled, 40 μL 3x firefly assay buffer was added and mixed vigorously to aid cell lysing. Plates were then incubated for 10 minutes while mixing at 400 rpm on a benchtop orbital shaker. After 10 minutes, the firefly luciferase activity was read using a Biotek Synergy plate reader with an integration time of 0.6 seconds. All luciferase reads were made twice in quick succession and later averaged before taking the ratio between luciferase activities. After the firefly signal was read, 60 μL of 3x nano luciferase assay buffer was added per well and mixed. The plates were then left to mix again for 10 minutes while shaking at 400 rpm. After 10 minutes, the second luciferase signal was measured. Firefly readouts were then normalized to the corresponding nano luciferase readout by dividing the firefly RLU by nano luciferase RLU for each individual well. Fold-changes were then calculated by averaging the luciferase ratio for the negative control and dividing all biological replications by the negative control’s luciferase ratio.

### Western blotting

48 hours after transfection for Neuro-2a experiments or 2 weeks after transduction for primary rat neuron experiments, media was removed from each well and the treated cells were washed with ice cold PBS and lysed in ice cold RIPA buffer (50 mM Tris, 150 mM NaCl, 1% Triton X-100, 0.5% sodium deoxycholate, 0.1% SDS, 1 mM EDTA, pH 7.4) supplemented with protease inhibitors and phosphatase inhibitors (Cell Signaling, 5872S). After 10 min incubation at room temperature with rocking, for experiments measuring Na_V_1.1 levels whole lysates were collected and frozen at −80C, but for experiments measuring CHD2 or ARID1B levels lysates were first centrifuged and the supernatant collected to remove debris. In Na_V_1.1 experiments, lysates were later thawed on ice and the DNA pellet was concentrated by light centrifugation and removed. 1 μL Pierce™ Universal Nuclease for Cell Lysis (Thermo Scientific) was then added to lower lysate viscosity and incubated on ice for at least 10 minutes.

Total protein concentration was measured by BCA assay (Thermo Scientific). The total protein amount loaded was confirmed to be within the linear range of detection for each antibody to detect each target protein for at least a 2x increase in protein amount (**Supplementary Fig. S7A and B**). For Na_V_1.1 experiments 16 μg (Neuro-2a) or 25 μg (primary rat neuron) total protein was mixed with 4X protein loading buffer (50 mM Tris pH 6.8, 2% SDS, 10% glycerol, 0.05% bromophenol blue, 100 mM DTT) and allowed to incubate at room temperature for 10 minutes. For CHD2 or ARID1B experiments 15 μg or 10 μg total protein respectively was mixed with 4X protein loading buffer and heated at 90 °C for 10 minutes. Prepared lysates were loaded on 8% SDS-PAGE gels and run at 100V during stacking and then 150V until the dye front reached the bottom of the gel.

The proteins were transferred onto a methanol activated PVDF membrane (pore size 0.45 µm; Immobilon-P from Millipore) using a wet transfer system (Bio-rad). Membranes were blocked with 3% BSA in TBST (200 mM Tris, 1500 mM NaCl, 0.01% Tween 20, pH 7.5) buffer for 1 h at room temperature, incubated with primary antibody (see **Supplementary Table S5** for antibodies and dilutions used in this study) in 3% BSA-TBST at 4 °C overnight. Membranes were then washed with TBST buffer three times for 10 minutes each, followed by corresponding HRP-conjugated secondary antibody in 3% BSA-TBST incubation 1 h at room temperature. The loading control α-tubulin was visualized using 1:5000 HRP-conjugated anti-α-tubulin antibody (Proteintech, HRP-66031). Membranes were imaged on a LI-COR Odyssey XF imager after incubation with SuperSignal West Pico PLUS chemiluminescent substrate (Thermo Scientific). Full western membrane images can be found in the **Source data file**.

When conducting the Na_V_1.1 western blot for the cortex and hippocampus tissue samples we saw that Na_V_1.1 presented as two bands, one at the correct 250 kDa size and one above it. Both bands seemed to be approximately half as intense in the SCN1a^+/-^ samples as the SCN1a^+/+^ samples leading us to believe that both bands represented different Na_V_1.1 folding variants or complexes since our sample processing steps were only mildly denaturing as to avoid unwanted aggregation. To highlight this fact, most examples of Na_V_1.1 blots in the literature include mildly heating the samples before loading, and when this is performed only one band appears at the incorrect, larger size though the correct correlation between genotype and band intensity is maintained (**Supplementary Fig. S7C**). Since our original western blot method faithfully reflects the correct protein size and sample genotype, we decided to move forward quantifying both bands together for future *in vivo* Na_V_1.1 western blot analysis.

### Quantification of signal intensity on western blots

All image analysis was performed in EmpiriaStudio. The chemiluminescence channel image was processed such that the box area drawn over each band was equal and individually centered on the band(s) intensity being measured. The mean intensity is then divided by the corresponding loading control intensity. The given intensity ratio is then normalized to the ratio of the negative control (set at 1) for final quantification and reporting. Quantification was only performed within target and loading control blot pairs and then combined across separate blot pairs for biological replicates to make western blot quantitation figures.

### RT-qPCR

For experiments in Neuro-2a cells, plating and transfection were performed as previously described. Total RNA was harvested 48 h after transfection and isolated using the RNeasy Mini Kit (QIAGEN). After isolation, RNA was reverse transcribed to cDNA using the PrimeScript RT Reagent Kit (TaKaRa). All qPCR reactions were run at 20 µL volumes in a 384-well plate (Applied Biosystems) with at least 3 biological replicates and 2 technical replicates for each biological replicate using PowerUp SYBR Master Mix (Thermo Scientific) and amplified on a QuantStudio 6 (Thermo Scientific). The qPCR primers were either identified based on previous publications or verified for specificity using NCBI Primer BLAST. Expression levels were calculated using the housekeeping control gene (GAPDH) cycle threshold (Ct) value and the gene of interest Ct value after averaging both technical replicates. The relative expression level of one gene was determined by 2^(−ΔCt), where ΔCt = Ct (gene of interest) − Ct (GAPDH). Relative expression level for targeted gene was obtained upon dividing the targeted gene expression level of cells experiments treated with the on-target CIRTS gRNA by those treated by the non-targeting (NT) CIRTS gRNA or vehicle for *in vivo* experiments. All qPCR primers can be found in **Supplementary Table S4**.

### Viral Production and Purification

For lentivirus production, HEK293T cells were plated on a 10 cm dish so that they would be approximately 70% confluent 24 hours later. The next day, media was replaced with 6 mL DMEM + 5% FBS and cells were transfected with four plasmids: VSV-G envelope plasmid (Addgene #12259), pRSV-Rev (Addgene #12253), pMDLg/pRRE (Addgene #12251), and lentiCRISPR v2 modified to replace CRISPR protein and gRNA cassettes with CIRTS equivalents described in a 2:2:2:4 μg ratio using 3 μg acidified polyethylenimine (PEI). 3 hrs after transfection, media was replaced with DMEM + 10% FBS. At 24 hrs after transfection, virus-containing media was collected and replaced. At 48 hrs after transfection, virus-containing media was again collected and combined with the media taken previously. The virus-containing media was then centrifuged at 500 g for 5 min and the supernatant was filtered through a 0.45 μm PES filter to remove cellular debris. The virus was then concentrated with PEG-it Virus Precipitation Solution (System Biosciences, LV810A-1) following manufacturer’s instructions and viral titer was estimated via qPCR via Lentivirus Titer Kit (Applied Biological Materials, LV900). Virus was then used immediately or frozen at −80C in single use aliquots.

For AAV production, CIRTS-4GT3/SCN1a-g1 was packaged into AAV9 by PackGene Biotech at a concentration of 1E13 GC/mL. Viral aliquots were kept at −80 °C and used immediately upon thawing.

### Primary neuronal culture and Lentiviral Transduction

Primary cultures of rat cortical neurons were prepared as described using Neurobasal Media (NBM), 4% (v/v) B27, and 0.125 mM l-glutamine (Thermo Fisher Scientific, Waltham, MA). Dissociated cortical neurons from E18 Sprague Dawley rat pups were plated in six-well plates coated with poly-D-lysine (Sigma, St Louis, MO) with 2 mL prepared media. For western blots, 0.4 × 10^6^ cells were plated in each well. At day 4 of culture, lentivirus at an MOI of 10 was diluted in 1 mL neuron media for each well to be transduced. 1 mL media was removed from the treated well and replaced with the media with virus. After transduction, 1 mL media was fed to the neurons approximately every 5 days. Transduction was confirmed for lentivirus encoding CIRTS fused with GFP by imaging by an inverted epifluorescence microscope (Leica DMi8) with a 10× objective, a Hamamatsu Orca-Flash 4.0 camera, and a 300 W Xenon light source (Sutter Lambda XL). Wells treated with lentivirus encoding CIRTS-4GT3 or CIRTS-NY1 with on-target or NT gRNA were processed as described in Western Blotting.

### Animals

Heterozygous SCN1a^+/-^ mice used as breeders to establish the colony used in this study, were 129S6/SvEvTac-Scn1a^tm1Kea^/Mmjax (RRID:MMRRC_037107-JAX) obtained from the Mutant Mouse Resource and Research Center (MMRRC) at The Jackson Laboratory, an NIH-funded strain repository, and was donated to the MMRRC by Jennifer Kearney, Ph.D., Northwestern University(36). The colony was propagated by crossing these SCN1a^+/-^ breeders with wildtype 129S6/SvEvTac mice (Taconic). For experiments, SCN1a^+/-^ mice on the 129S6/SvEvTac background were crossed with wild type C57BL/6N (Charles River) mice to generate F1 C57/126S9 offspring. Genotyping was performed via PCR as described by the MMRRC and Jackson Laboratory (see **Supplementary Table S4** for primer sequences).

All mice were maintained on a 12:12-hr light:dark cycle and had unrestricted access to food and water. Mice were euthanized using CO2 followed by cervical dislocation. For western blot analysis, cortical and hippocampal tissues were collected, combined and homogenized via a KIMBLE dounce tissue grinder (Sigma-Aldrich) in RIPA buffer + protease and phosphatase inhibitors prepared as described in Western Blotting. Samples were then sonicated for 5 sec at 25% amplitude and centrifuged at 14,000 g for 10 min at 4 °C. The supernatant was collected and used immediately for western blot as described previously or frozen at 80 °C. For RT-qPCR analysis, cortical and hippocampal tissues from one hemisphere were immediately placed in 500 µL RNAlater (Thermo Fisher Scientific) until RNA extraction by RNeasy Mini Kit (QIAGEN) following manufacturer’s instructions for RNA extraction from tissues. Downstream RT-qPCR was performed as previously described.

### i.c.v. injections

At postnatal day 1 (P1), neonates were anesthetized on ice for ∼3 min. 4 µL of viral suspension or vehicle control with added 0.05% Fast Green FCF dye (Sigma-Aldrich) to visualize successful intraventricular injection was injected into the right ventricle using a Model 701 RN 10 µL Syringe and 33-gauge, Small Hub RN Needle (Hamilton, 7635-01 and 7803-15) mounted on a micromanipulator (WPI: M3301R). The pup’s head was positioned exposing its right side, and the injection site was located by estimating roughly 1/2 the distance along an imaginary line between Lambda and the center of the eye. The needle was lowered to a depth of 2mm. Successful injection was confirmed by visualizing the characteristic winged shape of the dye representing the ventricles on either hemisphere. After injection, mice were placed on a warming pad and returned to the mother in the cage once recovered.

### Hyperthermia-induced Seizure (HTS) Assay

Mice were fitted with a rectal probe (RET-3; Physitemp) secured by medical tape and allowed to acclimate to the test chamber for 10 min. The probe was connected to a TCAT 2DF animal temperature controller (Physitemp) to continuously monitor the mouse’s internal body temperature. After 10 min, a heat lamp above the test chamber was turned on and the mouse’s body temperature was continuously monitored to increase by 0.5 °C every 2 min until onset of a tonic-clonic seizure, identified by erratic, uncontrolled movements and/or loss of posture, or a max internal body temperature of 42.5 °C was maintained for 3 min (as has been described previously (37,38)). Either temperature seizure first occurred, or the experiment was seizure free was recorded and mice were immediately placed in a recovery chamber with unrestricted food and water for at least 10 min.

### Statistics and reproducibility

All data are presented as mean ± SEM with individual data points. Statistical significance was assessed by unpaired two-tailed Student’s *t* test, one-way analysis of variance (ANOVA), two-way analysis of variance (ANOVA), or log-rank test with appropriate *post hoc* tests as described in the figure legends. P < 0.05 is considered statistically significant. Exact P values are provided in the source data file. All experiments were performed three or more times independently under identical or similar conditions, except when indicated in the figure legends.

## Results

### Identifying CIRTS-activator candidates for SCN1a translational activation

Previously, we found that a CIRTS fusion to the N-terminal domain (1-364) of the m6A reader protein YTHDF1 (CIRTS-NY1) was capable of activating translation from reporter genes and model transcripts (20,39). Therefore, we first assessed SCN1a activation using CIRTS-NY1, focusing on targeting the 3’ UTR of the SCN1a transcript, as we previously found targeting this region most effectively increased translation with CIRTS-NY1 (20). To facilitate screening and optimization, we constructed a dual luciferase reporter where the mouse 3’ UTR of SCN1a is expressed downstream of the Firefly Luciferase (Fluc) open reading frame, and an untargeted, destabilized Nano Luciferase (Nluc), is also expressed and is used as a transfection reference (**Fig. 2A**).

**Figure 2.**
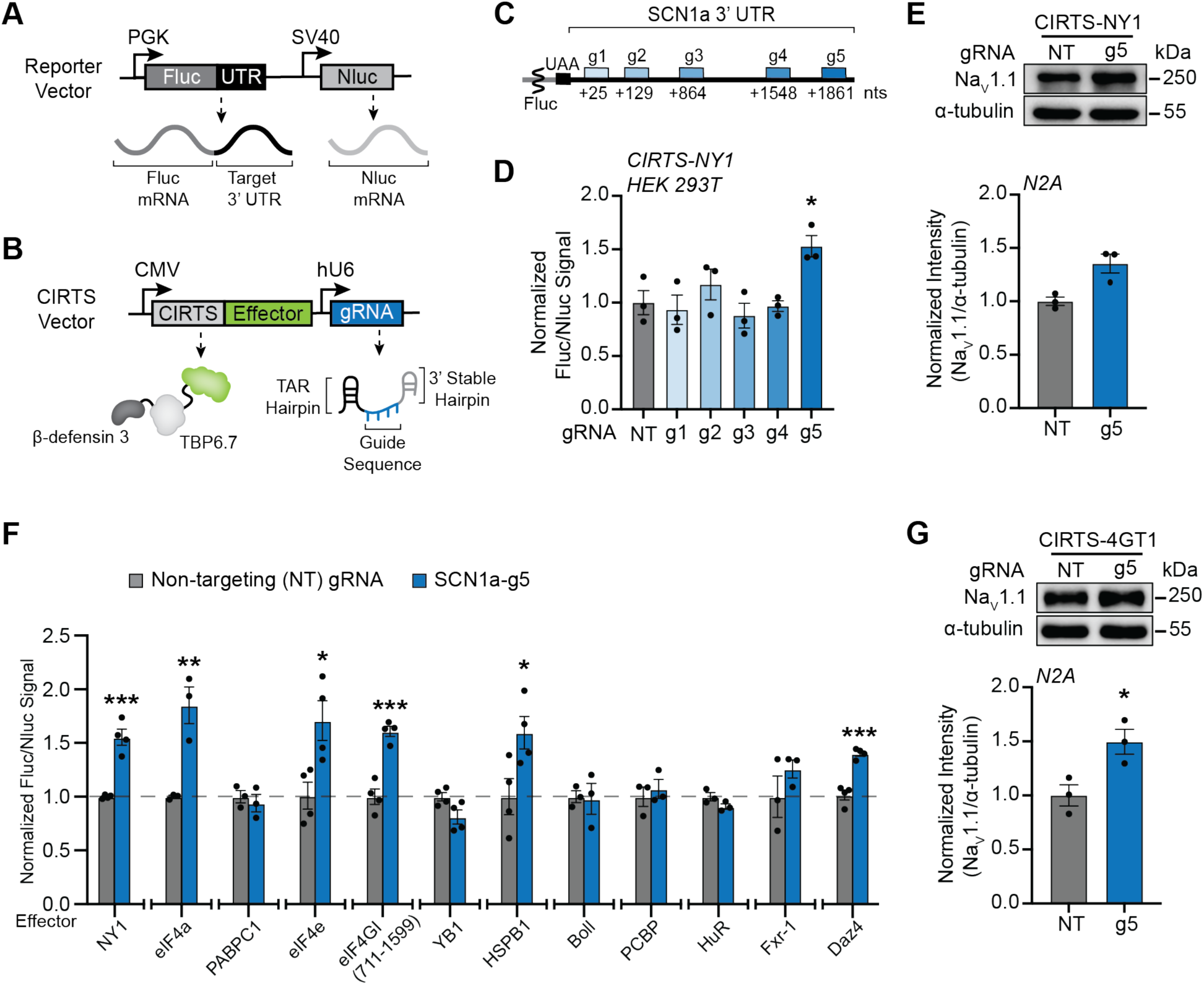
Screen for novel CIRTS-activators that increase endogenous Na_V_1.1 expression. (**A**) Vector design for the dual luciferase reporter, including expression cassettes for the Fluc with target 3’ UTR and the untargeted Nluc transcript used for transfection normalization. (**B**) Design for the combined CIRTS protein and gRNA expression vector used in this study. (**C**) Map of the SCN1a 3’ UTR, showing the distance in ribonucleotides (nts) from the start of the 3’ UTR to the first base to which each gRNA binds. (**D**) Evaluation of SCN1a 3’UTR-targeting CIRTS gRNA expressed alongside CIRTS-NY1 increasing Fluc expression by dual luciferase assay by two plasmid transfection in HEK293T cells. Each data point is normalized first to the internal Nluc transfection control and then against Fluc expression in the CIRTS-NY1 + non-targeting (NT) gRNA control. n = 3 biological replicates for each group. (**E**) Representative western blot for Na_V_1.1 levels 48 hours post-transfection with indicated CIRTS-NY1/gRNA vectors in N2a cells. α-tubulin was used as the loading control and SCN1a-g5 was normalized to NT gRNA for quantitation. n = 3 biological replicates for each group. (**F**) Screen for translational effectors fused with CIRTS targeting the SCN1a 3’ UTR dual luciferase reporter with SCN1a-g5. Each CIRTS was individually normalized to an effector-matched NT gRNA control. n = 3 or 4 biological replicates for each group. (**G**) Representative western blot for CIRTS-4GT1 expressed alongside SCN1a-g5 targeting endogenous SCN1a transcript in N2a cells 48 hours post-transfection. α-tubulin was used as the loading control and SCN1a-g5 was normalized to effector-matched CIRTS expressed with NT gRNA for quantitation. All bar graph values are shown as mean ± SEM with data points. Statistical analysis were performed using one-way ANOVA with *post-hoc* Dunnett’s multiple comparisons test (**D**) vs. NT. *\*P*<0.05. Unpaired two-tailed Student’s *t* test was performed in (**E**), (**F**), and (**G**) vs. effector-matched NT. *\*P*<0.05, *\*\*P*<0.01, *\*\*\*P*<0.001, *\*\*\*\*P*<0.0001. No asterisk = not significant.

We cloned dual expression vectors that express both CIRTS-NY1 (**Fig. 2B**) and a CIRTS gRNA, which we designed to be complementary to sites tiled along the SCN1a 3’ UTR (**Fig. 2C**). We then cotransfected each CIRTS/gRNA expression vector with the SCN1a 3’ UTR reporter into HEK293T cells and evaluated target Fluc activation normalized to transfection control. From this gRNA screen, we identified SCN1a-g5 as the optimal gRNA complimentary to the SCN1a 3’ UTR to mediate increased Fluc expression by CIRTS-NY1 compared to CIRTS-NY1 expression alongside a non-targeting (NT) gRNA (**Fig. 2D**). Next, we moved to confirm that the CIRTS-NY1/SCN1a-g5 pair can also increase translation from the endogenous SCN1a transcript by transfecting the expression vector into mouse Neuro-2a (N2a) cells, which natively express SCN1a. Western blot analysis revealed only modest activation of Na_V_1.1 levels compared to transfected CIRTS-NY1/NT vector (**Fig. 2E**, **Supplementary Fig. S2A**). While these results confirm CIRTS-NY1/SCN1a-g5 can target endogenous SCN1a and that SCN1a translation activation is feasible, they also indicate that the CIRTS-NY1 activator system is not likely to increase Na_V_1.1 protein to therapeutic levels. Because of the limited gRNA flexibility shown by CIRTS-NY1 on SCN1a, coupled with the complexity of further engineering NY1 due to the undefined mechanism by which it impacts translation (40,41), we were motivated to expand beyond the CIRTS-NY1 system for CIRTS-mediated translational activation. Therefore, we performed an effector screen, with the goal of identifying more robust and flexible SCN1a translational activators, including those amenable to optimization through protein and gRNA engineering.

We selected 11 candidate effector domains from the literature, primarily identified from “tethering” reporter assays, where the protein of interest was fused to an RNA-binding protein that interacts with multiple copies of a target RNA sequence on a reporter gene (42). We cloned CIRTS-effector domain fusions for each candidate into the dual expression vector with SCN1a-g5 and tested each construct using the SCN1a 3’ UTR luciferase reporter, using a NT gRNA as a control and CIRTS-NY1 as a reference. From this screen, we identified several effectors that mediate significant, gRNA-dependent increases in luciferase expression from the reporter (**Fig. 2F**). Putative new CIRTS activators include canonical eukaryotic initiation factors such as eIF4a and eIF4e, a C-terminal truncation of eIF4GI previously shown to drive translation in reporter assays, and HSPB1 and Daz4, which although having been previously identified in a high-throughput screen of human RBPs for translational regulators, have not yet been investigated in programmable RBP systems (43–45).

We next assessed whether each new CIRTS-activator can increase endogenous Na_V_1.1 expression targeting the native mRNA in N2a cells (**Supplementary Fig. S2B to F**). However, only CIRTS-eIF4GI (711–1599) (CIRTS-4GT1) demonstrated consistent and reproducible Na_V_1.1 expression (**Fig. 2G, Supplementary Fig. S2B**). While the expression increase imparted by CIRTS-4GT1 was modest and comparable to CIRTS-NY1, we selected CIRTS-4GT1 as our lead candidate to attempt to further optimize through gRNA screening and effector engineering, as the eIF4GI protein offers a high potential for further optimization due to greater characterization as a translational effector in the literature compared to YTHDF1 (46–49).

### Optimization of CIRTS-eIF4GI activators targeting SCN1a

To compare the flexibility of landing sites between effectors and to identify improved gRNAs, we tested CIRTS-4GT1 on the SCN1 3’ UTR reporter with the same gRNA panel we initially used for CIRTS-NY1. While only a single guide, SCN1a-g5, was capable of activation in the CIRTS-NY1 screen, both g1 and g4 showed significant activation when paired with CIRTS-4GT1 (**Fig. 3A**). Given the challenges observed from validating reporter results with the endogenous transcript, we next tested all three hit gRNAs with CIRTS-4GT1 on endogenous SCN1a, which revealed that all three guides increase Na_V_1.1 levels (**Fig. 3B**, **Supplementary Fig. S3A**), with g1 showing the highest activation, yielding a ∼50% increase in endogenous Na_V_1.1 expression compared to the NT gRNA.

**Figure 3.**
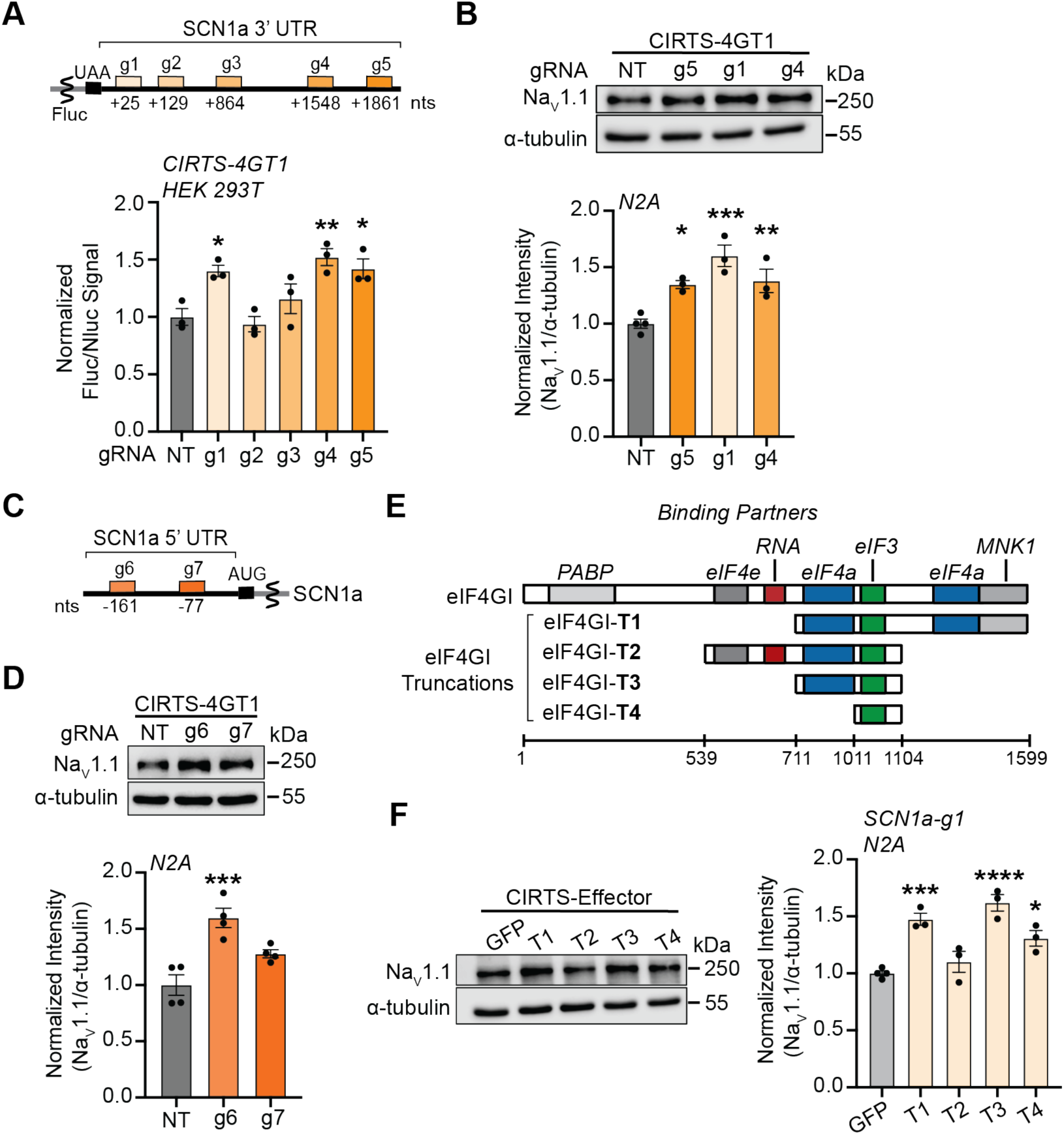
Optimizing CIRTS-eIF4GI through gRNA and protein engineering. (**A**) A SCN1a 3’ UTR gRNA panel was tested alongside CIRTS-4GT1 by dual luciferase assay in HEK293T cells. Relative Fluc expression was measured 48 hours post-transfection normalized to a transfected CIRTS-4GT1/NT gRNA control. n = 3 biological replicates. (**B**) Representative endogenous Na_V_1.1 western blot 48 hours post-transfection with vector expressing CIRTS-4GT1 and indicated gRNA. α-tubulin was used as the loading control and each gRNA was normalized to CIRTS-4GT1 expressed with NT gRNA for quantitation. n = 3 biological replicates. (**C**) SCN1a 5’ UTR map showing the distance in nucleotides (nts) from the start codon and the first nucleotide on the 5’ UTR each gRNA binds to. (**D**) Representative endogenous Na_V_1.1 western blot 48 hours post-transfection with vector expressing CIRTS-4GT1 and indicated 5’ UTR gRNA. α-tubulin was used as the loading control and each gRNA was normalized to CIRTS-4GT1 expressed with NT gRNA for quantitation. n = 4 biological replicates. (**E**) Map demonstrating eIF4GI’s domains and their binding partners over the full-length amino acid sequence. Tested truncations are shown below the full-length protein with the domains they possess highlighted. (**F**) Representative endogenous Na_V_1.1 western blot 48 hours post-transfection with vector expressing the indicated CIRTS-effector and SCN1a-g1 α-tubulin was used as the loading control and each CIRTS-4G truncation was normalized to CIRTS-GFP, where GFP was fused to CIRTS as the effector domain, expressed with SCN1a-g1. n = 3 or 4 biological replicates. All bar graph values are shown as mean ± SEM with data points. Statistical analyses were performed using one-way ANOVA with *post hoc* Dunnett’s multiple comparisons test vs. NT (**A**), (**B**), and (**C**) or vs. GFP (**E**) *\*P*<0.05, *\*\*P*<0.01, *\*\*\*P*<0.001, *\*\*\*\*P*<0.0001. No asterisk = not significant.

We next aimed to further map the generality of landing sites for CIRTS-4GT1. eIF4GI is an important member of the eIF4F complex (along with eIF4e and eIF4a), which is integral for scaffolding translation initiation and is thought to canonically act in the 5’ UTR (50). We therefore tested whether CIRTS-4GT1 can also promote translation of SCN1a from the 5’ UTR. Because the 5’ UTR of SCN1a is much shorter than its 3’ UTR (187 nts compared to 2103 nts), we designed only two gRNAs, g6 and g7, targeting the 5’ UTR, and tested them directly by Western blot on endogenous SCN1a in N2a cells. We found that g6 could direct CIRTS-4GT1 to increase Na_V_1.1 expression compared to a NT gRNA (**Fig. 3C**, **Supplementary Fig. S3B**). Of note, we found both g6 and g7 failed to activate Na_V_1.1 expression with both CIRTS-NY1 and -eIF4e (**Supplementary Fig. S3C and D**), further confirming the increased landing site flexibility offered by CIRTS-4GT1 compared to CIRTS-NY1 and CIRTS-4GT1. The amenability of CIRTS-4GT1 for targeting both 5’-and 3’-UTRs would likely be beneficial when developing potent, specific translational activators for new endogenous targets.

With improved gRNAs targeting both the 3’ and 5’ UTRs (g1 and g6, respectively), we next sought to identify the minimal protein domains required for the 4GT1 effector to increase protein expression. eIF4GI is an important protein for scaffolding and orchestrating initiation factors in the early stages of translation and is composed of many protein-binding domains (**Fig. 3D**) (46). Previously, tethering assays used to investigate eIF4GI truncations both *in vitro* and in cells demonstrated that the central core of the protein, spanning the middle eIF4a binding domain and eIF3 binding domain, is essential for increasing translation from reporters (44,47–49). We hypothesized that the eIF3 interaction is likely critical for activity, as eIF4a itself was a non-functional CIRTS effector. Moreover, we recently demonstrated eIF3 recruitment by RNA-based IRES elements in *trans* also functions as a viable translational activator (51).

We designed three additional eIF4GI truncations, each retaining the core eIF3 binding domain, and probed how each domain outside of the eIF3 binding domain contributes to increasing Na_V_1.1 protein expression as a CIRTS-effector (**Fig. 3D**). eIF4GI-T2 lacks the C-terminal eIF4A and Mnk1 domains, but adds in the eIF4e binding domain and the RNA binding domain (RBD) upstream of this functional core, which, in *in vitro* tethering assays (49), improved translation stimulation in 5’ UTR. eIF4GI-T3 only includes the middle eIF4a and eIF3 binding domains, and eIF4GI-T4 includes only the eIF3 binding domain.

We tested each CIRTS-delivered effector on endogenous SCN1a activation using the 3’ UTR-targeting g1 (Fig. 3E, Supplementary Fig. S3E) and 5’ UTR-targeting g5 (**Supplementary Fig. S3F and G**), to assess how effectors act on the transcript in different contexts. These experiments revealed that the C-terminal eIF4a and Mnk1 binding domains are not major contributors to effector activity in the 3’ UTR, in agreement with previous tethering assays with the 5’ UTR (44). Surprisingly, including the eIF4e and RBD on the effector resulted in subpar activation when directed to the 3’ UTR, and removing the middle eIF4a domain dramatically decreased activity, supporting *in vitro* evidence that eIF4a and eIF3 may bind cooperatively to this eIF4GI region creating a bridge between the bound mRNA and 40S ribosome independently of the RBD (49,52). Results in the 5’ UTR also confirmed the C-terminal region after amino acid 1104 does not contribute significantly to activity, but strikingly, including the eIF4e and the RBD was beneficial in this context. Furthermore, deleting the middle eIF4a binding domain also negatively impacted effector function in the 5’ UTR, further confirming that the core eIF4a and eIF3 binding domains together are the main drivers of effector activity.

From these results, we selected the CIRTS-eIF4G-T3 (CIRTS-4GT3) effector for subsequent studies, as it demonstrated the most robust and general activity across a variety of target locations. Additionally, we anticipate removal of the promiscuous RBD should decrease effector binding to nonspecific RNAs, thus greatly improving specificity, as the CIRTS-4GT3 fusion should therefore depend solely on the gRNA for mRNA-engagement (53). We also prioritized further development with CIRTS-4GT3 directed by 3’ UTR SCN1a-g1, as targeting the 3’ UTR is likely more versatile as a general gRNA design strategy for future target-development – human 3’ UTRs are, on average, longer, less structured, and alternatively spliced to a lesser extent than 5’ UTRs (54).

### CIRTS-4GT3 is an improved translational activator and deployable by lentivirus in primary neurons

To benchmark the performance of CIRTS-4GT3 with CIRTS-NY1, we compared each optimized activator/gRNA pair targeting SCN1a in N2a cells by Western blot (**Fig. 4A**, **Supplementary Fig. S4A**) and RT-qPCR analysis (**Fig. 4B**). We observed that the CIRTS-4GT3/SCN1a-g1 pair drives a 90% increase in expression, outperforming CIRTS-NY1/SCN1a-g5, which averaged a 50% increase. The approximate doubling of Na_V_1.1 expression observed with CIRTS-4GT3/SCN1a-g1 suggests it is a compelling candidate for correcting Na_V_1.1 under-expression in haploinsufficiency without risking toxicity from protein overexpression. Importantly, CIRTS-4GT3 does not significantly impact target transcript levels during expression activation, confirming that the increase in Na_V_1.1 is driven by translational activation, in agreement with previous characterization of the CIRTS-NY1 mechanism (20,40). Furthermore, a slight decrease in mRNA level is observed, consistent with decreased mRNA stability during heavy translation, and which is not recapitulated upon on-target gRNA binding with CIRTS-GFP (55).

**Figure 4.**
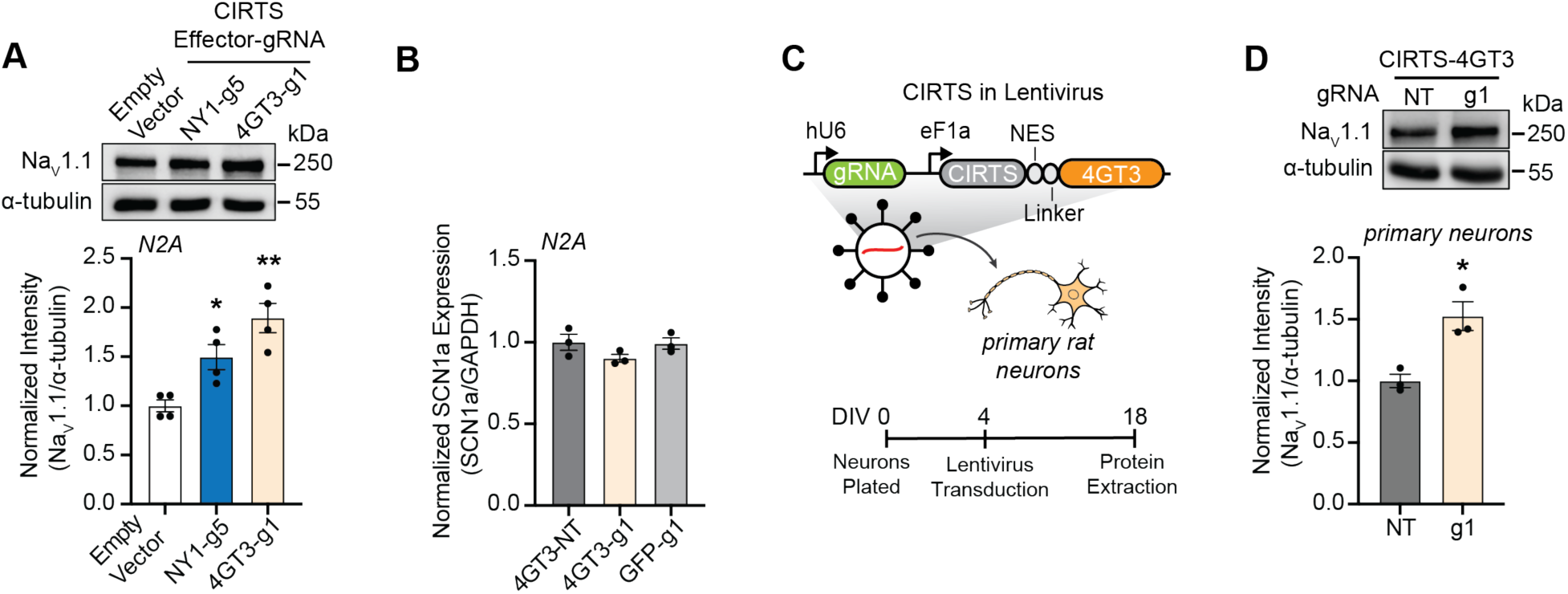
CIRTS-4GT3 is an improved CIRTS-activator and active in primary neurons. (**A**) Vectors expressing either CIRTS-NY1/SCN1a-g5, CIRTS-4GT3/SCN1a-g1 or empty CIRTS/gRNA cassettes were transfected into N2a cells and endogenous Na_V_1.1 levels were measured 48 hours later. Representative western image is shown. α-tubulin was used as the loading control and both groups were normalized to the empty vector for quantitation. n = 4 biological replicates (**B**) SCN1a mRNA levels were measured by RT-qPCR 48 hours after transfection with vectors expressing either CIRTS-4GT3 or CIRTS-GFP and either on-target SCN1a-g1 or NT gRNA. GAPDH was used as the reference gene and each group was normalized to the CIRTS-4GT3/NT condition. n = 3 biological replicates. (**C**) Vector map for the CIRTS-4GT3/gRNA lentivirus expression cassettes and experimental design for primary rat cortical neuron transduction and protein extraction. (**D**) Representative western blot analysis of Na_V_1.1 levels in primary cortical rat neurons 2 weeks post-transduction with lentivirus expressing CIRTS-4GT3/NT or CIRTS-4GT3/SCN1a-g1 at an MOI = 10. α-tubulin was used as the loading control and signal was normalized to CIRTS-4GT3 expressed with NT gRNA for quantitation. n = 3 biological replicates. All bar graph values are shown as mean ± SEM with data points. Statistical analyses were performed using one-way ANOVA with *post hoc* Dunnett’s multiple comparisons test vs. empty vector (**A**), or with *post hoc* Sidak’s multiple comparisons test vs. CIRTS-4GT3/NT gRNA (**B**). Statistical analyses were performed using unpaired two-tailed Student’s *t* test vs. NT (**D**). *\*P*<0.05, *\*\*P*<0.01. No asterisk = not significant.

Next, we aimed to evaluate the functionality of CIRTS-4GT3 in a neuronal context and test whether SCN1a translation activation is sustained when CIRTS-4GT3 and on-target gRNA are constitutively expressed in rat primary cortical neurons after viral delivery. We designed a CIRTS lentiviral vector based on previously reported CRISPR/gRNA expression systems (56) (**Fig. 4C**). To confirm CIRTS expression and optimize delivery, we first transduced primary rat cortical neurons at 4 days *in vitro* (4 DIV) with a CIRTS-GFP/NT-gRNA lentiviral vector. Fluorescence microscopy at 10 DIV showed that a multiplicity of infection (MOI) of 10 results in approximately 100% transduction (**Supplementary Fig. S4B and C**).

After confirming CIRTS-GFP is stably expressed in primary neurons, we generated lentiviral vectors expressing CIRTS-4GT3 and either SCN1a-g1 or NT-gRNA. SCN1a-g1 was fully complementary to the rat SCN1a 3’ UTR and the sequence was unmodified compared to past experiments. We transduced neurons on 4 DIV with one of the two lentiviral systems at 10 MOI and then measured Na_V_1.1 levels by Western blot analysis at 18 DIV (**Fig. 4C**). As expected, neurons treated with lentivirus encoding on-target gRNA expressed approximately 50% more Na_V_1.1 than neurons transduced with CIRTS-4GT3/NT gRNA, confirming that CIRTS-4GT3 functions in a primary neuron context, with measurable effects at 2 weeks post-transduction (**Fig. 4D**, **Supplementary Fig. S4D**). Despite the worse performance of CIRTS-NY1 in preliminary assays, because YTHDF1 has been shown to actively regulate translation in neurons (57), we also tested CIRTS-NY1/SCN1a-g5 delivered by lentivirus transduction to rat primary cortical neurons. Indeed, these experiments revealed a significant gRNA-dependent increase in Na_V_1.1 expression at 2 weeks-post transduction (**Supplementary Fig. S4E to G**), revealing that CIRTS-NY1 is also a viable translational activator in primary neurons, though CIRTS-4GT3 was still prioritized for future experiments due to its higher activation efficiency.

### CIRTS-4GT3 improves phenotype in a Dravet syndrome mouse model

With evidence that virally delivered CIRTS-4GT3 targeting SCN1a mRNA has the potential to drive sustained Na_V_1.1 levels in primary neurons, we next sought to determine if CIRTS-4GT3 could boost Na_V_1.1 expression *in vivo* and improve phenotype in a SCN1a haploinsufficiency model. Heterozygous knockout of SCN1a in mice (SCN1a^+/-^) faithfully recapitulates many aspects of Dravet Syndrome (DS) in human patient populations (58). These symptoms include marked early life mortality primarily due to sudden death in epilepsy (SUDEP) and a susceptibility to hyperthermic-induced seizures (HTS), which corroborates the 15-20% mortality in DS patients before 20 years of age and fever being a common trigger for DS-related epileptic episodes (36). Crucially, DS mouse models are also very sensitive to Na_V_1.1 levels; previous studies demonstrate triggered re-expression or viral delivery of SCN1a rescues DS phenotypes, with even a 25% boost in expression contributing to measurable phenotypic improvements (59). These aspects make SCN1a^+/-^ mice a valuable testbed for validating CIRTS-activators as a viable strategy for addressing haploinsufficiency on both a molecular and phenotypic level.

Previous work in DS mouse models characterized GABAergic inhibitory interneurons in the hippocampus and cortex as a crucial cell type impaired by reduced Na_V_1.1 expression and key driver of the epileptic phenotype (60). These cells have been the choice target for DS genetic-targeting therapies in order to best balance phenotypic improvements with mitigating off-target effects; however, Na_V_1.1 is also expressed in other neuronal cell types with undetermined contribution to the DS phenotype that are likely missed by interneuron-specific delivery and expression strategies (27–29). We sought to overcome this limitation and leverage the intrinsic selectivity for cells expressing SCN1a transcript afforded by our mRNA-targeting approach and designed a CIRTS-based therapy for broad neuronal delivery. To this end, we designed and generated an AAV9 expressing CIRTS-4GT3 driven by a pan-neuronal hSYN promoter and SCN1a-g1 under a constitutive hU6 promoter (AAV9-4GT3-g1) (**Fig. 5A**).

**Figure 5.**
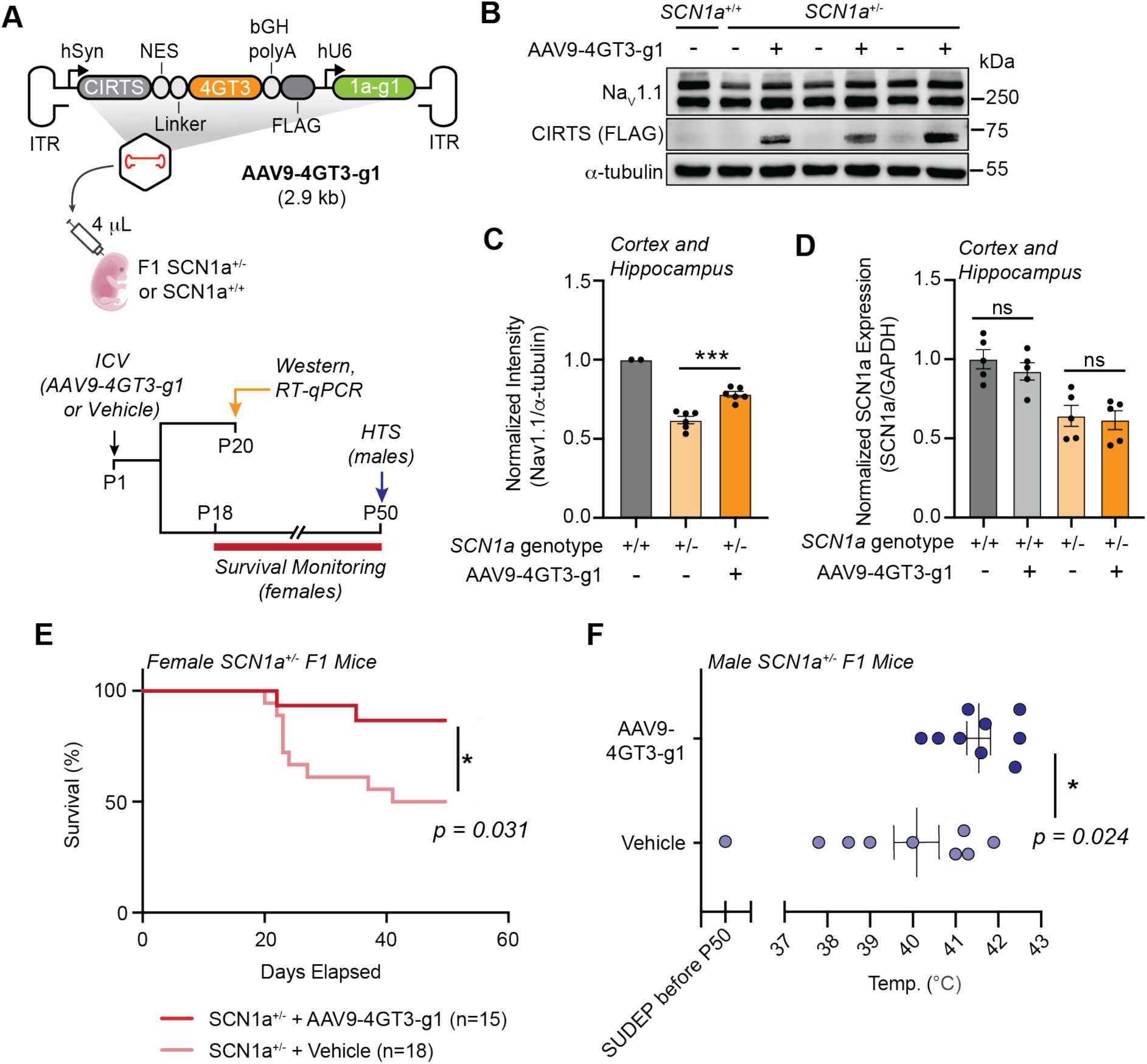
CIRTS-4GT3 in AAV9 delivered by i.c.v. injection to a Dravet syndrome mouse model increases SCN1a expression and improves phenotype. (**A**) CIRTS-4GT3/SCN1a-g1 vector map for neuron-specific CIRTS-4GT3 expression via AAV9 delivery and experimental schematic for mouse cohorts used to measure indicated molecular and phenotypic endpoints. (**B**) Representative western blot assay of Na_V_1.1 levels in P20 female F1 mice cortex and hippocampus tissue samples. Genotype for each sample is indicated above western blot as well as treatment with either i.c.v. injection on P1 of AAV9-4GT3-g1 or vehicle. Western blot analysis for P20 male F1 mice can be found at **Fig. S6A** α-tubulin was used as the loading control. (**C**) Quantitation for western blot analysis shown in Fig. 5B and **Fig. S6A** Normalization was carried out in reference to Na_V_1.1 levels in wt F1 mice treated with vehicle. n = 2 for wt mice or = 6 biological replicates for SCN1a^+/-^ mice. (**D**) SCN1a mRNA levels measured in both wt and SCN1a^+/-^ P20 male F1 mice cortex and hippocampus tissue samples after treatment with either AAV9-4GT3-g1 or vehicle via i.c.v. injection on P1. GAPDH was used as the reference gene. n = 5 biological replicates (**E**) Survival rate in female SCN1a^+/-^ F1 mice treated with either AAV9-4GT3-g1 or vehicle via P1 i.c.v. injection after symptoms onset at ∼P18 to P50. (**F**) HTS assay performed on male P50 F1 SCN1a^+/-^ mice treated with either AAV9-4GT3-g1 or vehicle via P1 i.c.v. injection. One male mouse treated with vehicle did not survive to P50 and was not included in statistical analysis. n = 9 biological replicates. All bar and scatter plot graph values are shown as mean ± SEM with data points. Statistical analyses were performed using two-way ANOVA with Sidak’s multiple comparisons test between vehicle- and AAV9-4GT3-g1-treated SCN1a^+/-^ mice (**C**) and between vehicle- and AAV9-4GT3-g1-treated SCN1a^+/+^ and SCN1a^+/-^ mice (**D**). Statistical analyses were performed using Log-rank test (**E**) between vehicle- and AAV9-4GT3-g1-treated female SCN1a^+/-^ mice and unpaired two-tailed Student’s *t* test (**F**) between vehicle- and AAV9-4GT3-g1-treated male SCN1a^+/-^ mice. *\*P*<0.05, *\*\*P*<0.01, *\*\*\*P*<0.001.

Genetic background is important for DS mice to develop the epileptic phenotype.

Previous work has established SCN1a^+/-^ mice have no overt phenotype on a 129S6/SvEvTac background, but if crossed with C57BL/6J their F1 offspring (129S6/SvEvTac x C57BL/6J) will exhibit symptom onset at approximately P21 and experience approximately 50% mortality with death by SUDEP typically occurring in the third and fourth weeks of life.

In order to evaluate the viability of this strategy to correct SCN1a^+/-^ phenotype in these F1 offspring, we first sought to test whether AAV9-4GT3-g1 increases Na_V_1.1 expression in the cerebral cortex and hippocampus by P21. To ensure broad neuronal AAV transduction and CIRTS-4GT3 expression by this time point we choose to deliver AAV9-4GT3-g1 by neonatal intracerebroventricular (i.c.v.) injection as this method has been demonstrated to be effective for cerebral and hippocampal AAV9-mediated transgene expression within P21 previously (61).

Western blot analysis of cortico-hippocampal samples from F1 P21 mice confirmed specific CIRTS expression in animals treated with AAV9-4GT3-g1 compared to animals treated with vehicle, demonstrating sustained CIRTS-4GT3 expression in the brain (**Fig. 5B, Supplementary Fig. S5A**). Correspondingly, Na_V_1.1 protein levels were modestly increased (∼30%) in AAV9-CIRTS-treated mice compared to control (**Fig. 5C, Supplementary Fig. S5A**). As expected, SCN1a mRNA levels were not significantly affected by AAV9-4GT3-g1 treatment in either genotype, further confirming the increase in Na_V_1.1 levels *in vivo* were likely due to increased translation (**Fig. 5D**). This improvement produces Na_V_1.1 levels below that of wildtype; however, given that even modest increases in Na_V_1.1 have been shown to measurably ameliorate DS-like symptoms, we were nevertheless encouraged to pursue phenotypic studies to further characterize the potential of SCN1a-targeting CIRTS-4GT3 as a novel therapeutic modality. We therefore tested if the Na_V_1.1 expression boost measured resulted in functional protein capable of impacting the epileptic phenotype.

A second F1 cohort was treated with AAV-CIRTS or vehicle, and were monitored for survival until P50, to assess the impact AAV9-4GT3-g1 treatment had on early-life mortality in SCN1a^+/-^ mice. Among SCN1a^+/-^ mice, we observed a dramatic difference in mortality between control males and females, with control female mice displaying the commonly reported 50% mortality for this model: 9 out of 18 female mice, as compared to only 1 out of 9 males dropping out before P50 (**Fig. 5E and Supplementary Fig. S5B**). While sexual dimorphisms in disease presentation are not uncommon in epilepsies, there has been conflicting evidence for the extent this DS model presents these differences (36,62,63). Due to the phenotypic differences observed in this study, we evaluated male and female mice separately for all subsequent phenotypic evaluation, in agreement with several recent publications using this model (62–64). Remarkably, for female SCN1a^+/-^ mice, AAV9-4GT3-g1 treatment resulted in a significantly reduced mortality rate of 13% (n = 2/15), compared to 50% for vehicle-treated female SCN1a^+/-^ mice survival until P50 (n = 9/18) (**Fig. 5E**). We did not observe any deaths in the female or male SCN1a^+/+^ AAV9-CIRTS or vehicle treatment groups, underscoring the efficacy and safety potential for this strategy (**Supplementary Fig. S5B and C)**.

Next, we evaluated if AAV9-4GT3-g1 treatment impacted HTS susceptibility in SCN1a^+/-^ mice as the temperature threshold at which seizures are observed is another important measurable phenotype for DS severity (23). After only one vehicle-treated male SCN1a^+/-^ dropped out in the F1 survival cohort, we tested the remaining mice by HTS assay in order to compare the temperature threshold for an induced seizure at P50 between treatment groups.

We found SCN1a^+/-^ mice treated with AAV9-4GT3-g1 possessed a significantly higher temperature threshold (41.5 °C) for induced seizures compared to mice treated with vehicle (40.1 °C) (**Fig. 5F**). Additionally, we recorded no seizures in SCN1a^+/+^ mice from either treatment group (**Supplementary Fig. S5D)**. Altogether these results confirm that SCN1a-targeting CIRTS-4GT3 improves functional, neuronal Na_V_1.1 expression *in vivo* and has exciting potential as a preclinical modality to address DS.

### CIRTS-4GT3 can programmably activate translation for other haploinsufficiency targets

Inspired by the preclinical success CIRTS-4GT3 demonstrated in the DS mouse model we sought to assess whether CIRTS-4GT3 could also improve the translation of other endogenous transcripts with validated haploinsufficiency-related disorders. To this end we selected two genes, *CHD2* and *ARID1B*, which encode proteins that contribute to chromatin remodeling in neurons that are too large for traditional gene replacement. *CHD2* haploinsufficiency results in epilepsy and developmental delay while *ARID1B* haploinsufficiency has been shown to cause intellectual disability and autism-spectrum disorder with no approved treatments available to correct protein under-expression for either condition (30,31).

Following a similar pipeline established for SCN1a gRNA screening, we first constructed a dual luciferase reporter with the mouse CHD2 3’ UTR downstream of the Fluc CDS and designed 5 gRNAs tiling along the 3’ UTR (**Fig. 6A**). Plasmid encoding CIRTS-4GT3 and either NT or CHD2-targeting gRNA were transfected alongside the CHD2 3’ UTR dual luciferase reporter in order to identify functional CIRTS-4GT3/gRNA pairings (**Fig. 6B**). We observed CHD2-g2 mediates a significant increase in normalized Fluc expression with CIRTS-4GT3 with CHD2-g3 and CHD2-g4 showing potential, albeit unsignificant activity. We next moved to assess if our putative screening hit, CIRTS-4GT3/CHD2-g2, could activate endogenous CHD2 translation by transfection in N2a cells and western blot analysis (**Fig. 6C, Supplementary Fig. S6A**). Compared to CIRTS-4GT3/NT, CIRTS-4GT3/CHD2-g2 led to a significant (50%) increase to CHD2 expression, while CIRTS-4GT3/CHD2-g3 only weakly impacted CHD2 expression, in agreement with the luciferase assay. Additionally, we confirmed that activation with CIRTS-4GT3/CHD2-g2 does not significantly change CHD2 mRNA levels in N2a cells, and again we observed a slight decrease in mRNA stability upon targeting supporting CIRTS-4GT3 increases CHD2 translation **(Supplementary Fig. S4B**). Interestingly, we also repeated the CHD2 gRNA screen with CIRTS-NY1 and found that while CIRTS-NY1/CHD2-g2 and CHD2-g3 both significantly increased Fluc expression in the luciferase assay neither pair demonstrated an increase in endogenous CHD2 expression by western blot, further confirming CIRTS-4GT3 is an improved, and more general CIRTS-translational activator (**Supplementary Fig. S6C to F**).

**Figure 6.**
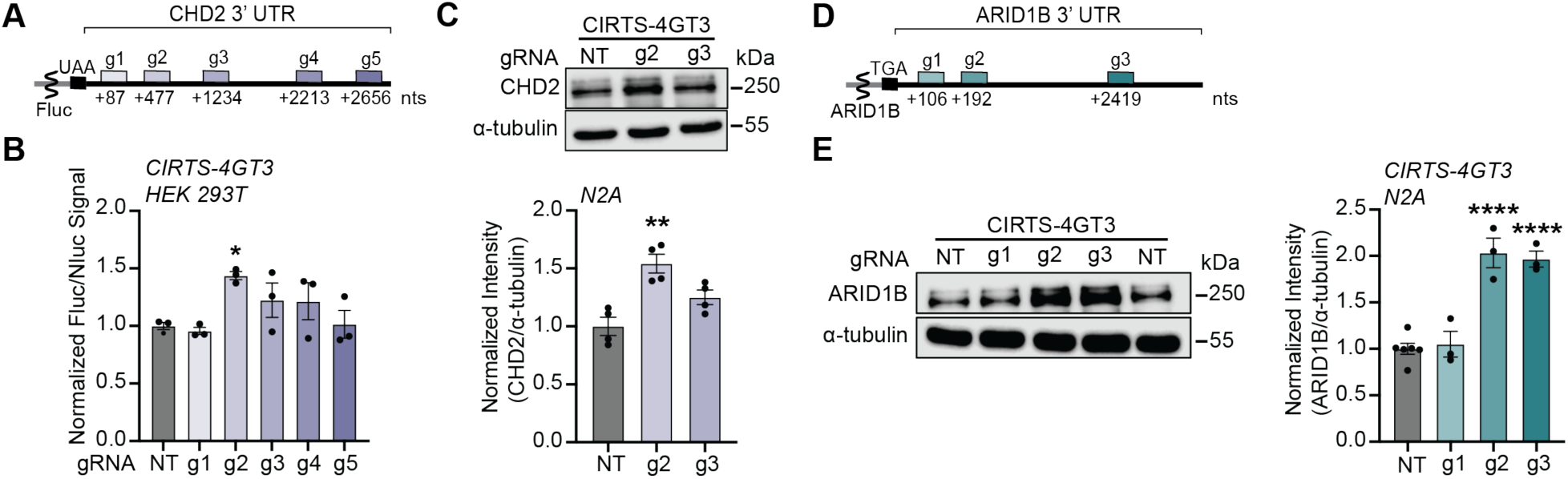
CIRTS-4GT3 can programmably activate translation for additional haploinsufficiency-related targets. (**A**) 3’ UTR map for a mouse CHD2 dual luciferase reporter with gRNA positions annotated showing the distance in nucleotides (nts) from the start of the 3’ UTR and the first nucleotide on the 3’ UTR each gRNA binds to. (**B**) CHD2 dual luciferase assay results 48 hours after HEK293T cells were transfected with vectors expressing CIRTS-4GT3 and a CHD2 gRNA panel member. Each replicate was divided by their respective Nluc value and normalized to their effector-matched, NT gRNA control. n = 3 biological replicates. (**C**) Representative endogenous CHD2 western blot completed with N2a cell lysate collected 48 hours post-transfection with vectors expressing CIRTS-4GT3 and indicated gRNA. α-tubulin was used as the loading control and each gRNA was normalized to effector-matched CIRTS expressed with NT gRNA for quantitation. n = 4 biological replicates. (**D**) 3’ UTR map for the mouse ARID1B 3’ UTR with gRNA positions annotated showing the distance in nucleotides (nts) from the start of the 3’ UTR and the first nucleotide on the 3’ UTR each gRNA binds to. (**E**) Representative endogenous ARID1B western blot completed with N2a cell lysate collected 48 hours post-transfection with vectors expressing CIRTS-4GT3 and indicated gRNA. α-tubulin was used as the loading control and each gRNA was normalized to effector-matched CIRTS expressed with NT gRNA for quantitation. n = 6 or 3 biological replicates. Statistical analyses were performed using one-way ANOVA with *post hoc* Dunnett’s multiple comparisons test vs. CIRTS-4GT3/NT gRNA (**B**), (**C**), and (**E**). *\*P*<0.05, *\*\*P*<0.01, *\*\*\*P*<0.001, *\*\*\*\*P*<0.0001. No asterisk = not significant

While the luciferase assay is helpful and accurate for identifying lead gRNA for a new target, cloning out full 3’ UTRs from cellular RNA can be time consuming. Given we had quickly found hits in each designed gRNA group for both SCN1a and CHD2, for identifying CIRTS-4GT3/gRNA pairs that increase ARID1B translation we designed a smaller panel of three gRNA and tested them directly with CIRTS-4GT3 by transfection in N2a cells and subsequent western blot analysis (**Fig. 6D and 6E**). ARID1B western blot revealed that both ARID1B-g2 and ARID1B-g3 produced significant (100%) increases to ARID1B expression compared to cells transfected with CIRTS-4GT3/NT. Overall, these results demonstrate that CIRTS-4GT3 is a general, programmable translational activator that likely has the potential to correct protein under-expression in haploinsufficiency beyond Na_V_1.1 and is primed for future preclinical development.

## Discussion

In this work, we optimized CIRTS-4GT3, a 601 aa protein that can activate translation from three different haploinsufficiency-related mRNAs, is easily deliverable by therapeutically tractable single viral vectors and, to our knowledge, represents the first translational activator system shown to positively impact a haploinsufficiency phenotype *in vivo.* For DS specifically, CIRTS-4GT3 joins an exciting panel of emerging preclinical and early clinical therapeutics aimed at alleviating SCN1a haploinsufficiency, while possessing a unique attribute blend in the field (65). Current strategies to address SCN1a haploinsufficiency act primarily on either the DNA-level or RNA-level. Genomic technologies include gene supplementation using dual-AAVs or adenovirus due to *SCN1a*’s large size, or gene activation with CRISPR/dCas9 or Zinc finger transcriptional activator fusions (27,66–69). RNA-targeting has also been demonstrated to be effective via ASO-based technologies that either improve productive SCN1a mRNA splicing, or knockdown a repressive lncRNA (59,70). CIRTS-4GT3 uniquely combines several advantages from both groups, specifically, transcriptome-level specificity of RNA-targeting and the ability to be genetically encoded and delivered broadly by AAV for sustained Na_V_1.1 expression rescue in one dose. The advantages of transcriptome-level specificity is especially apparent when comparing the increased mortality reported when using the pan-neuron hSYN1 promoter to express Na_V_1.1 itself in a similar DS mouse model with the improvement to survival reported in this study when expressing CIRTS-4GT3 instead from the same promoter (27). However, future work will be needed to investigate neuron population-specific SCN1a activation from AAV9-4GT3-g1 transduction and whether pan-neuron or GABAergic inhibitory interneuron-specific expression of CIRTS-4GT3 has differing impact on disease phenotype.

While this work supports the ability of translational activation to address SCN1a haploinsufficiency in a heterozygous knockout DS model, many diverse SCN1a mutations have been identified in patient populations. The majority of cases, and on average more severe, involve mutations which lead to no or truncated Na_V_1.1 expression, which will likely be amenable to translational activation (58). Future studies will be needed to examine how translational activation impacts DS derived from SCN1a missense mutations as a pool of mutant RNA would also be present and likely not cleared by nonsense mediated decay as in many truncation variants (71). If unwanted targeting of the mutant mRNA diminished activation of the healthy mRNA, then modalities such as shRNA or orthogonal CIRTS-degraders could potentially be included in the AAV vector to achieve mutant-selective knockdown to bias the targetable mRNA population. Furthermore, much like other developed SCN1a haploinsufficiency therapies, translational activation would not be amenable to other rare SCN1a-related DEEs derived from gain-of-function mutations which demonstrate dominant-negative pathology (25).

The phenotypic improvements measured in this study imparts vital preclinical support to the burgeoning translational activation field and advances its development as an impactful strategy to boost protein expression. Over the last decade, programmable translational activation in mammalian cells has made large strides and various programmable translational activation technologies have been developed (72,73). Current protein-based translational activation platforms function primarily through fusion of translation-activating effector proteins with RNA-binding peptide repeats such as engineered PUF or PPR domains or with gRNA-dependent CRISPR-based proteins (39,74,75). CIRTS has clinical advantages due to its compact size (dCas13d-4GT3 would be 1,391 aa, and a 30-nt targeting PPR-4GT3 would be ∼1,444 aa, both over the AAV packaging limit if inserted into the vector design used in this study) and its human origin, potentially mitigating the immunogenicity concerns associated with microbial-derived CRISPR-based technologies, which likely interfere with long-term treatments needed for haploinsufficiency (21,22). For effectors, eIF4GI and its domain truncations have emerged as a leading choice for translational-activation as they have been recently demonstrated to also be effective at boosting translation from endogenous mRNA with both dCas13d- and PPR-based fusions targeting the 5’ UTR (75,76). This current work contributes new evidence that eIF4GI truncations are also functional in activating translational when targeted to the 3’ UTR as well as the minimal domains required to drive this effect. The findings in this work highlight eIF4GI as a potent translational activator and motivates future efforts elucidating by what mechanism eIF4GI domain delivery to endogenous mRNA coordinates increased translation, especially in the 3’ UTR, as crosslinking between eIF4GI’s binding partner eIF3 and highly translated mRNA’s 3’ UTRs has been recently reported suggesting similar pathways may be natively active (77,78).

Oligo-based translational activator technologies which combine an RNA translation-activating domain including SINE B (SINEUPs) or IRES domains (taRNAs) directly with a gRNA also offer a potential avenue to increase target translation (51,79–81). However, SINEUPs are restricted to gRNA landing sites overlapping with the start codon, which may limit target flexibility depending on accessibility within this region or unavoidable sequence similarity with off-target transcripts. taRNAs, like CIRTS, offer greater gRNA design flexibility as they generally function by landing in either the 5’ and 3’ UTR, but have not demonstrated to function *in vivo* beyond LNP-mediated delivery to the liver and likely require further development to achieve sustained *in vivo* activity. Overall, boosting protein expression will likely not be “one size fits all” and a variety of therapeutic modalities with different attributes will be necessary to address haploinsufficiency broadly.

Beyond DS, CIRTS-4GT3 also has the potential to be applied to improve protein expression in other haploinsufficiencies. The pipeline demonstrated for gRNA design and screening by dual luciferase assay could allow for the rapid development of CIRTS-4GT3/gRNA pairs for a broad panel of targets. Interestingly, while the correlation between CIRTS-4GT3 function in the dual luciferase assay and on endogenous transcripts was strong, we found that the results from dual luciferase screening did not always translate to endogenous targeting for CIRTS-NY1 and the other translational activator domains tested. Since many studies to identify translational effectors in the literature depend on similar, often less stringent, high-throughput luciferase assays these results highlight the need for orthogonal, follow-up assays to confirm effectors can modify translation on endogenous transcripts. This observation also highlights the potential – and challenges – of targeting RNA regulation for therapeutic purposes, as nuances in cell-specific contexts can be critical for ultimate efficacy.

In sum, this work validates a new protein-based translational activation modality with flexible, programmable targeting with proven efficacy *in vivo* on a high-value therapeutic target. This work confirms that targeting translational activation has therapeutic value, at least in a *SCN1a* heterozygous knockout DS model, indicating the more work on translational activation technology development is warranted. The CIRTS platform is well positioned to offer a strategy for gene expression control in situations where gene replacement may be difficult to implement, or DNA-changes may be undesirable. Future work further improving the platform’s efficacy will likely yield potent preclinical candidates to address the great need for therapeutics to address haploinsufficiency as a general disease mechanism.

## Data availability

The data underlying this article are available in the article and in its online supplementary material or will be shared on reasonable request to the corresponding author.

## Supplementary data statement

Supplementary Data are available at NAR Online

## Acknowledgements

We thank Dr. Somayeh Ahmadiantehrani for editing assistance with the figures and manuscript, and Dr. Yang Cao, Dr. Tong Lan, and the rest of the Dickinson lab for helpful discussions during this work.

## Author contributions

Riley Sinnott (Conceptualization [lead], Data curation [lead], Formal analysis [lead], Investigation [lead], Methodology [lead], Visualization [lead], Writing-original draft [lead], Writing-review and editing [lead]), Ani Solanki (Investigation [supporting], Methodology [supporting]), Anitha P. Govind (Resources [supporting]), William N. Green (Funding acquisition [supporting]), Bryan C. Dickinson (Conceptualization [lead], Formal analysis [supporting], Funding acquisition [lead], Methodology [supporting], Supervision [lead], Writing-original draft [supporting], Writing-review and editing [supporting])

