## Supplementary Data for "Engineering a human-based translational activator for targeted protein expression restoration"

**TABLE OF CONTENTS**

**Supplementary Figure 1**: Overview on technologies to increase gene product

**Supplementary Figure 2**: Additional $Na_V$1.1 western blots for dual luciferase screening hits

**Supplementary Figure 3**: Extended western blots targeting SCN1a 5' and 3' UTRs with CIRTS-NY1, -eIF4e, and -eIF4GI truncations

**Supplementary Figure 4**: Primary rat neuron lentivirus transduction with CIRTS-GFP/NT and CIRTS-NY1/rSCN1a-g5 $Na_V$1.1 western blots

**Supplementary Figure 5**: Extended *In vivo* F1 DS mice western blot, survival, and HTS data

**Supplementary Figure 6**: Extended CHD2 and ARID1B western blots and RT-qPCR

**Supplementary Figure 7**: Linear detection range for western blot primary antibodies and *in vivo* $Na_V$1.1 western blot optimization

**Supplementary Table 1**: Representative plasmids used in this study

**Supplementary Table 2:** CIRTS gRNA sequences

**Supplementary Table 3:** CIRTS effectors amino acid sequences

**Supplementary Table 4:** Primer sequences

**Supplementary Table 5:** Antibodies and dilutions used for western blots

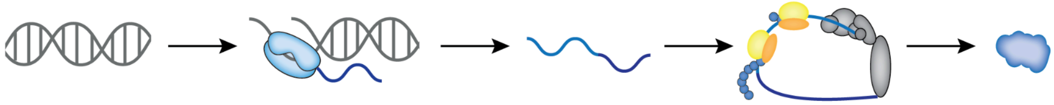

| Central Dogma: | DNA | Transcription | mRNA | Translation | Protein |
|---|---|---|---|---|---|
| **Technology:** | • Gene supplementation<br>• CRISPR/Cas9<br>• Base editing<br>• Prime editing | • CRISPRa<br>• Zinc finger (ZF) activators<br>• saRNA | • ASOs<br>• mRNA delivery<br>• RNA editing | • Suppressor tRNAs<br>• taRNAs<br>• SINEUPs<br>• CIRTS or dCas13d | • Protein delivery<br>• DUBTACs<br>• small molecule activators |
| **Advantages:** | • Potential for one treatment to correct disease<br>• Editors could correct disease even if delivered transiently<br>• Mutant-selective editing could address dominant negative disease-causing varients | • Potential for one treatment to correct disease (CRISPR or ZF activators)<br>• Can treat variety of mutations with single activator<br>• ZF activators fit into single AAV vectors and may be less immunogenic than CRISPRa | • RNA-delivery provides dosability and reversibility<br>• RNA cheaper to produce than viral vectors<br>• RNA-targeting provides transcriptome specificity<br>• Can treat variety of mutations with single ASO | • RNA-targeting provides transcriptome specificity<br>• Potential for one treatment to correct disease (if AAV is used for delivery)<br>• Can treat variety of mutations with single activator or targets with single tRNA | • Small molecules are relatively cheap to produce<br>• Provides dosability and reversibility |
| **Caveats:** | • Viral delivery limited by gene or editor size<br>• Cell-type delivery must be strictly controlled<br>• CRISPR immunogenicty<br>• Editors need to be transient to limit permanent off-target effects | • Viral delivery limited by activator size (CRISPRa)<br>• Cell-type delivery must be strictly controlled<br>• CRISPR or dsRNA immunogenicty<br>• Limited brain distribution via RNA-delivery (saRNA) | • Limited brain distribution via RNA-delivery<br>• Dependence on target regulatory mechanisms (ASOs)<br>• Limited by gene size (mRNA delivery) | • Limited brain distribution via RNA-delivery (t/taRNAs)<br>• Non-specific readthrough (tRNAs)<br>• Limited target flexibility (SINEUPs)<br>• Unoptimized efficiacy (tRNAs and CIRTS) | • Limited brain distribution (protein delivery)<br>• Must cross blood-brain-barrier<br>• Difficult to discover *de novo* (small molecules) |

**Supplementary Fig. 1.** Overview on technologies to increase gene product

Schematic representing the central dogma for genetic information flow from DNA to mRNA to protein is shown above. Technologies that can theoretically lead to increased correct gene product in haploinsufficency are listed with advantages and caveats under the molecule or transitional step in the central dogma they most directly influence.

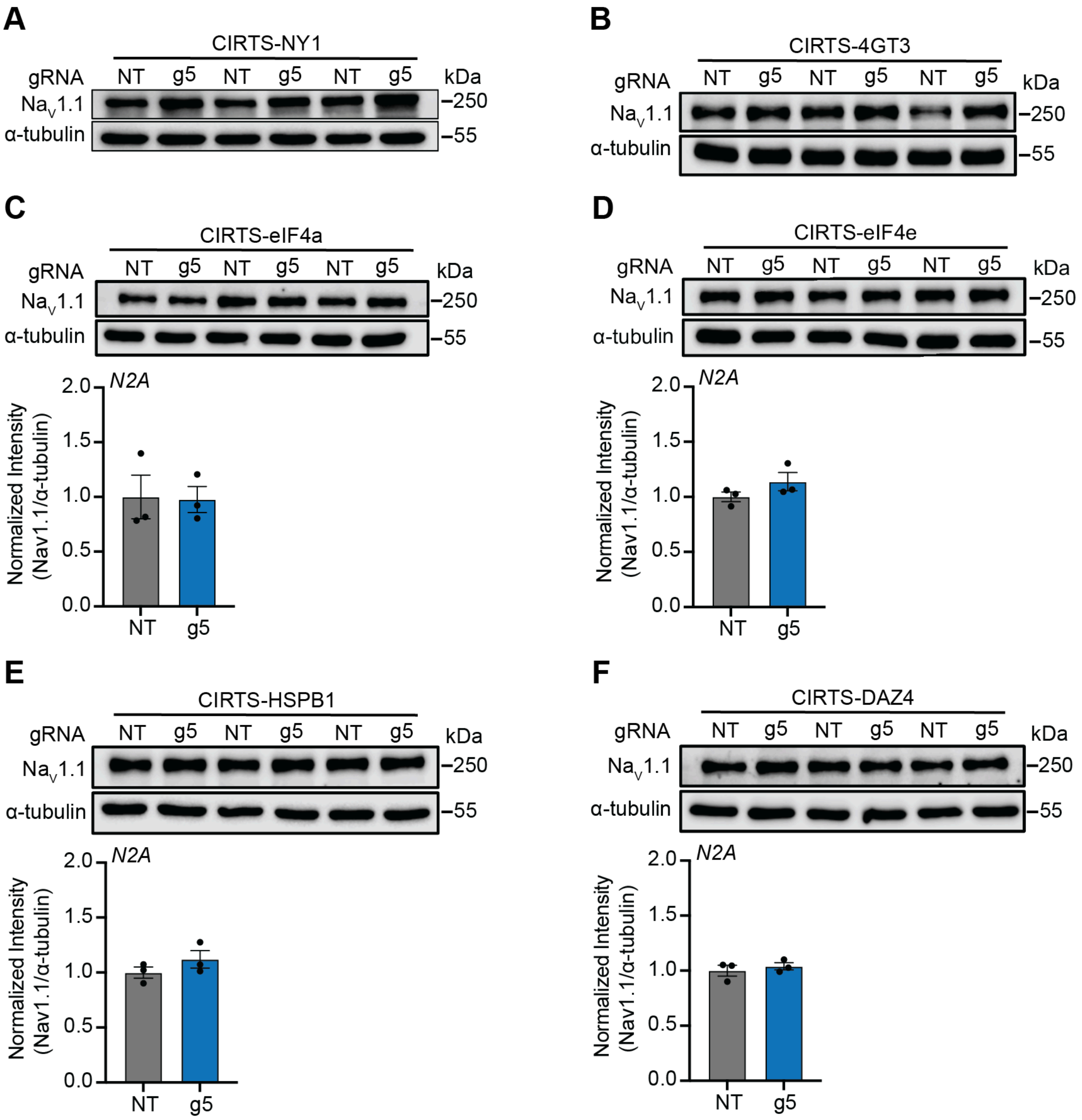


**Supplementary Fig. 2.** Additional $Na_V1.1$ western blots for dual luciferase screening hits

(**A**) Full western blot used in **Fig. 2E** quantitation. (**B**) Full western blot used in **Fig. 2G** quantitation. (**C**-**F**) western blots for $Na_V1.1$ levels 48 hours post-transfection with indicated CIRTS-effector/gRNA vectors in N2a cells. α-tubulin was used as the loading control and each CIRTS-effector/SCN1a-g5 was normalized to the matched CIRTS-effector expressed with NT gRNA for quantitation. n = 3 biological replicates for each group. All bar graph values are shown as mean ± SEM with data points. Statistical analyses were performed using unpaired two-tailed Student's *t* test vs. matched CIRTS-effector/NT (**C**-**F**). No asterisk = not significant.

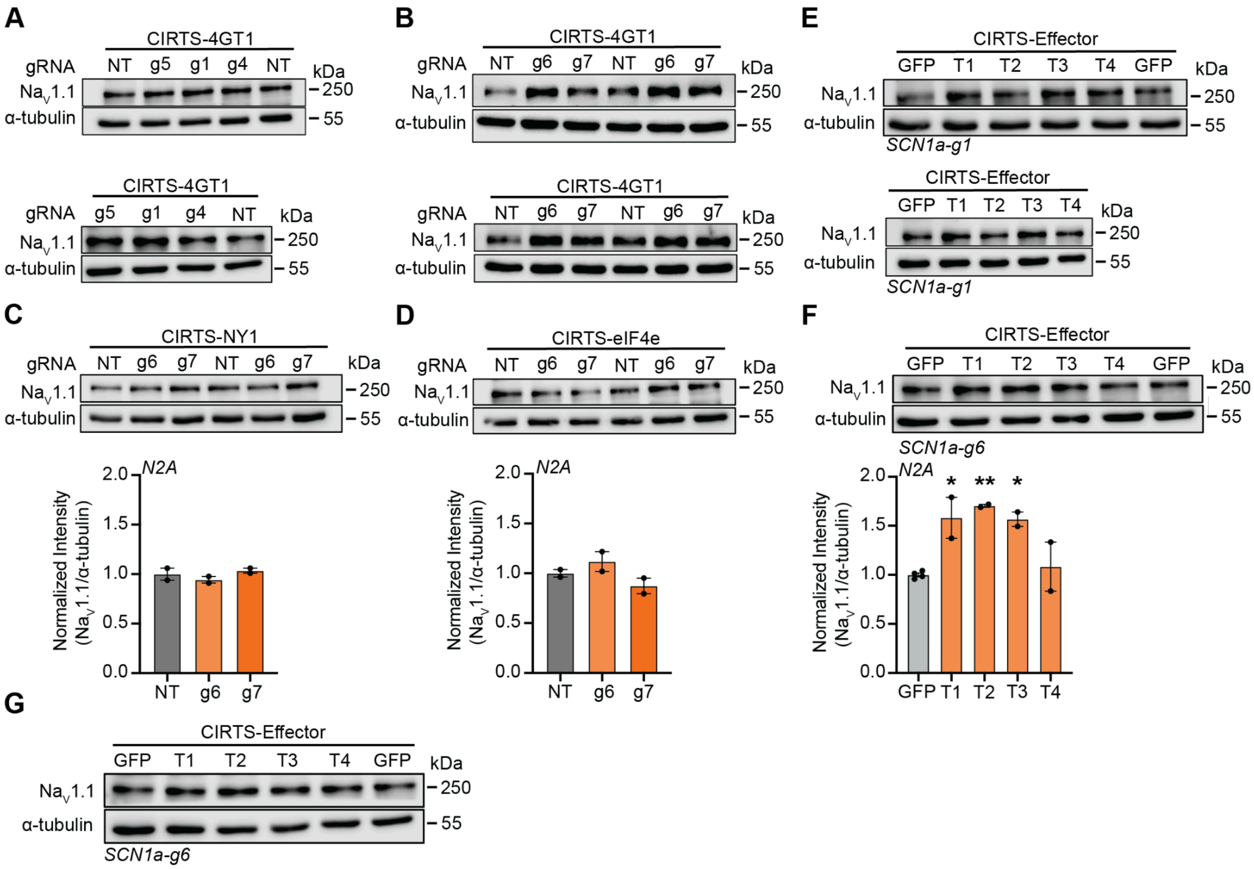


**Supplementary Fig. 3.** Extended western blots targeting SCN1a 5’ and 3’ UTRs with CIRTS-NY1, eIF4e, and -eIF4GI truncations

(**A**) Full western blots used in **Fig. 3B** quantitation. (**B**) Full western blots used in **Fig. 3D** quantitation. **(C-D)** Endogenous $Na_V$1.1 western blots 48 hours post-transfection with vectors expressing CIRTS-NY1 (**C**) or CIRTS-eIF4e (**D**) and indicated 5’ UTR-targeting gRNA. α-tubulin was used as the loading control and each gRNA was normalized to the matched CIRTS-effector expressed with NT gRNA for quantitation. n = 2 biological replicates. (**E**) Full western blots used in **Fig. 3F** quantitation. (**F**) $Na_V$1.1 western blot 48 hours post-transfection with vectors expressing the indicated CIRTS-effector and SCN1a-g6. α-tubulin was used as the loading control and each CIRTS-4G truncation was normalized to CIRTS-GFP/SCN1a-g6, n = 2 or 4 biological replicates. (**G**) Additional western blot used in **Fig. S3F** quantitation. All bar graph values are shown as mean ± SEM with data points. Statistical analyses were performed using one-way ANOVA with Dunnett’s multiple comparisons test vs. NT (**C**) and (**D**) or vs. GFP (**F**) *$P$<0.05, **$P$<0.01. No asterisk = not significant.

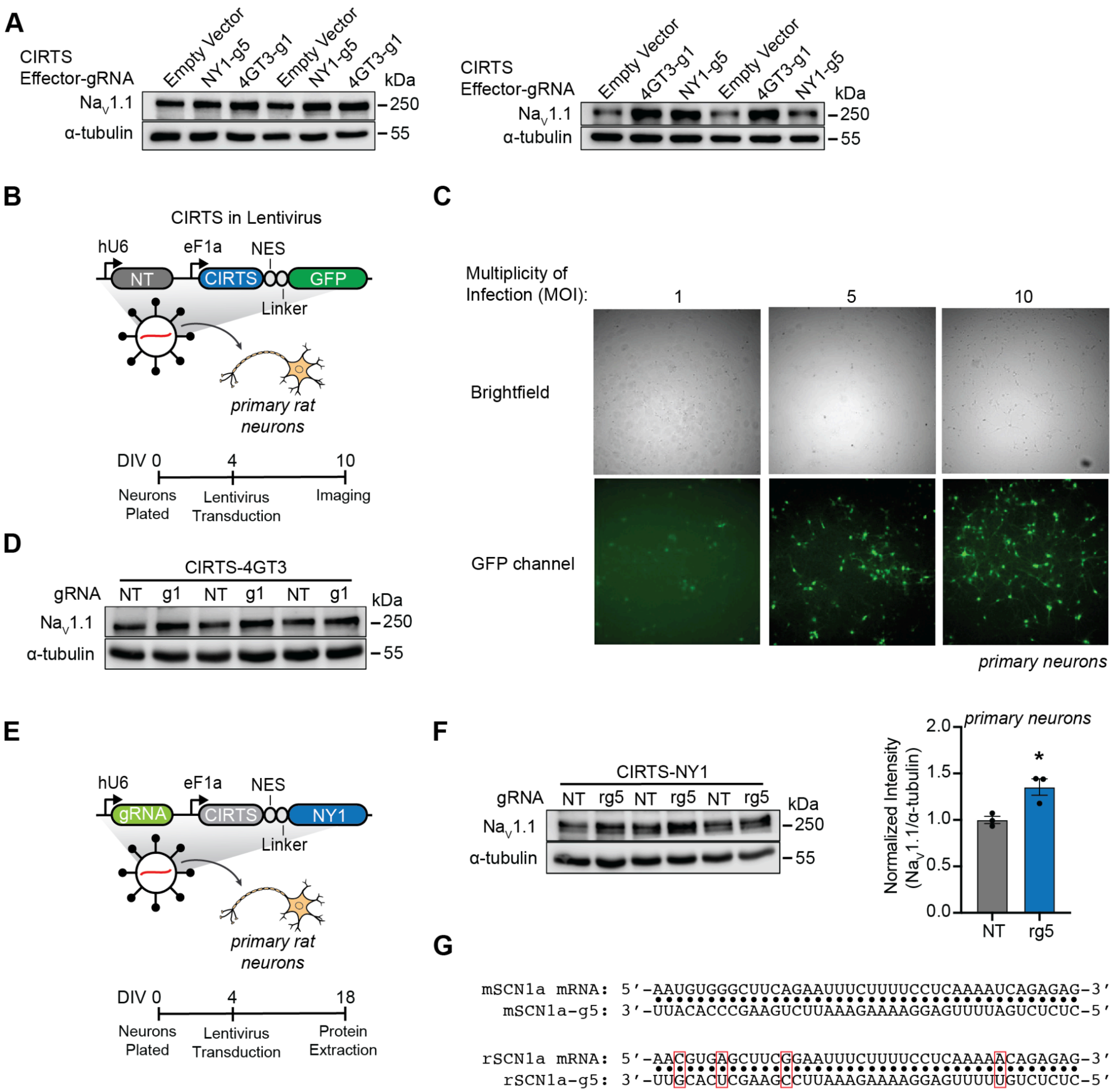


**Supplementary Figure 4**: Primary rat cortical neuron lentivirus transduction validation with CIRTS-GFP/NT and CIRTS-NY1/rSCN1a-g5 $Na_V$1.1 western blots

(**A**) Full western blots used in **Fig. 4A** quantitation. (**B)** Vector map for the CIRTS-GFP/NT gRNA lentivirus expression cassettes and experimental design for primary rat neuron transduction and GFP imaging. (**C**) Representation images of primary rat cortical neurons in the brightfield and GFP channel 6 days post-transduction with different MOIs of CIRTS-GFP/NT lentivirus. Images were captured and processed using the same settings between treated groups. (**D**) Full western blot used in **Fig. 4D** quantitation. (**E**) Vector map for the CIRTS-NY1/gRNA lentivirus expression cassettes and experimental design for primary rat neuron transduction and protein extraction. (**F**) Western blot analysis of $Na_V$1.1 levels in primary cortical rat neurons 2 weeks post-transduction with lentivirus expressing CIRTS-NY1/NT or CIRTS-NY1/rSCN1a-g5 at a MOI = 10. α-tubulin was used as the loading control and signal was normalized to CIRTS-NY1 expressed with NT gRNA for quantitation. n = 3 biological replicates. (**G**)

Alignment of partial sequences from mouse and rat SCN1a mRNA 3’ UTR with the mouse-targeting SCN1a-g5 gRNA and rat-targeting SCN1a-rg5 gRNA respectively. Bases modified in the SCN1a-rg5 sequence compared to the SCN1a-g5 sequence to ensure full target-complementarity are highlighted in red boxes. All bar graph values are shown as mean ± SEM with data points. Statistical analysis was performed using unpaired two-tailed Student’s *t* test vs. NT (**F**). **P*<0.05. No asterisk = not significant.

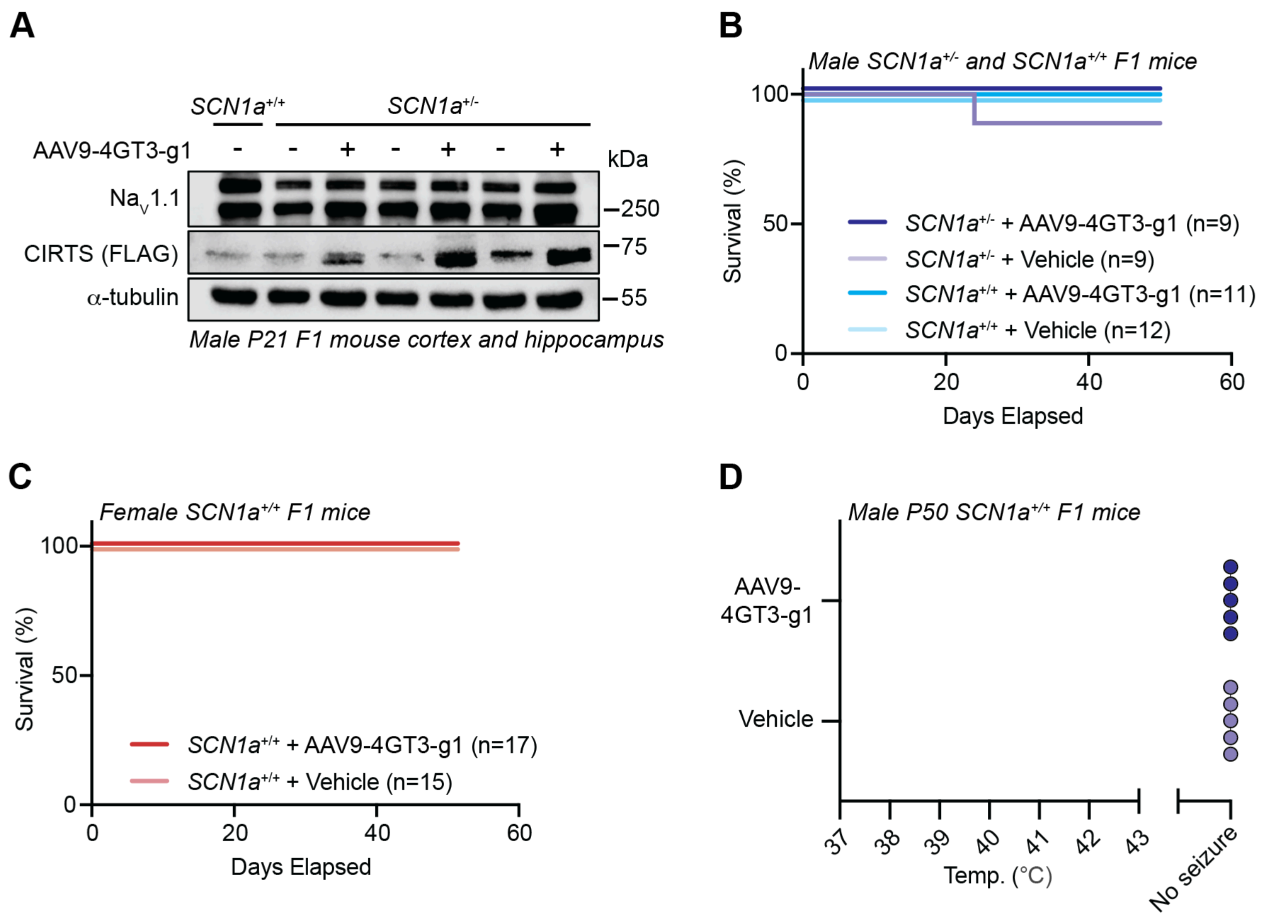


**Supplementary Figure 5**: Extended *In vivo* F1 DS mice western blot, survival, and HTS data

(**A**) Additional western blot used in **Fig. 5B** quantitation performed with cortex and hippocampus tissue from male P21 F1 mice treated with either AAV9-4GT3-g1 or vehicle via P1 i.c.v. injection. (**B**) Survival rate in male SCN1a$^{+/-}$ and SCN1a$^{+/+}$ F1 mice treated with either AAV9-4GT3-g1 or vehicle via P1 i.c.v. injection to P50. (**C**) Survival rate in female SCN1a$^{+/+}$ F1 mice treated with either AAV9-4GT3-g1 or vehicle via P1 i.c.v. injection to P50. (**D**) HTS assay performed on male P50 F1 SCN1a$^{+/+}$ mice treated with either AAV9-4GT3-g1 or vehicle via P1 icv injection. n = 5 biological replicates. All scatter plot graph values are shown as mean ± SEM with data points. Statistical analyses were performed using Log-rank test between all groups (**B**) and (**C**). No asterisk = not significant.

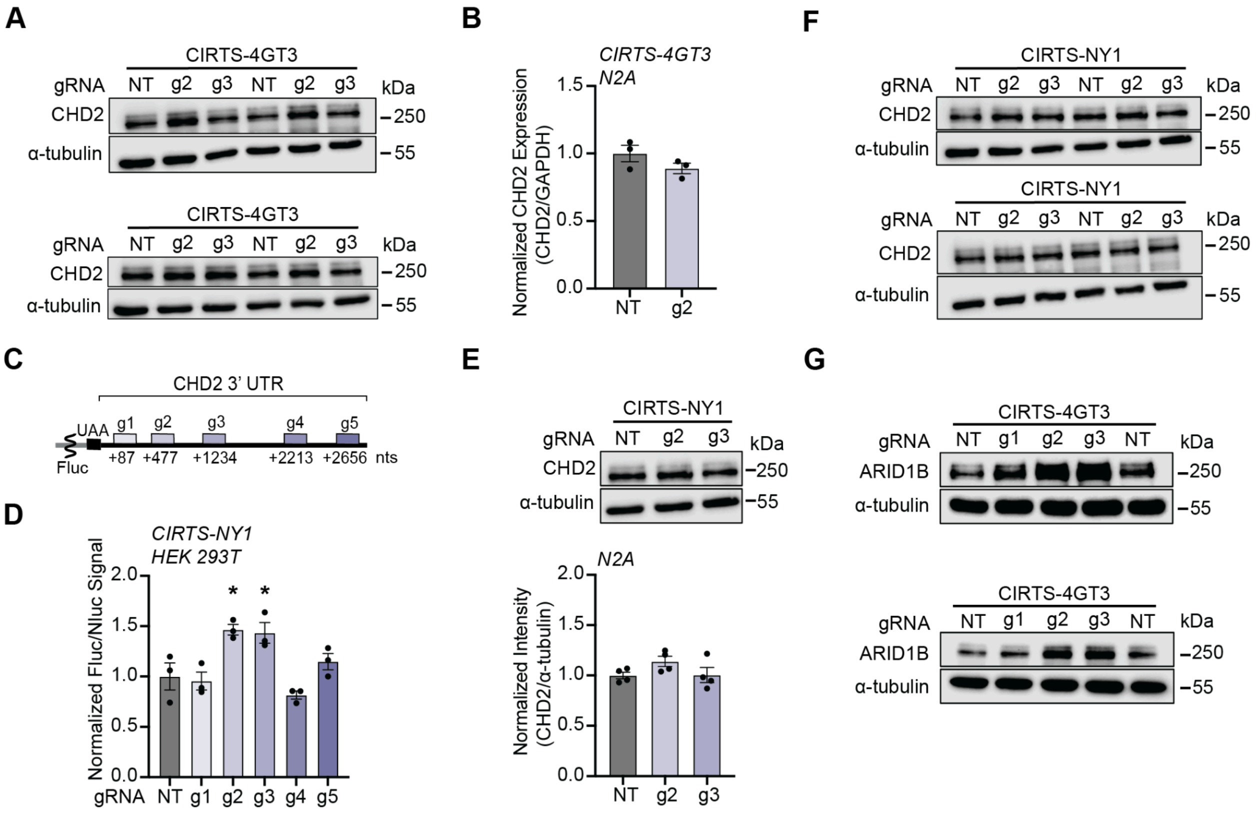


**Supplementary Figure 6**: Extended CHD2 and ARID1B western blots and RT-qPCR

(**A**) Full western blots used in **Fig. 6C** quantitation. (**B**) CHD2 mRNA levels were measured by RT-qPCR 48 hours after transfection with vectors expressing CIRTS-4GT3 and either on-target CHD2-g2 or NT gRNA. GAPDH was used as the reference gene and data was normalized to the CIRTS-4GT3/NT condition. n = 3 biological replicates. (**C**) 3' UTR map for a mouse CHD2 dual luciferase reporter with gRNA positions annotated showing the distance in nucleotides (nts) from the start of the 3' UTR and the first nucleotide on the 3' UTR each gRNA binds to. (**D**) CHD2 dual luciferase assay results 48 hours after HEK293T cells were transfected with vectors expressing CIRTS-NY1 and a CHD2 gRNA panel member. Each replicate was divided by their respective Nluc value and normalized to their effector-matched, NT gRNA control. n = 3 biological replicates. (**E**) Representative endogenous CHD2 western blot completed with N2a cell lysate collected 48 hours post-transfection with vectors expressing CIRTS-NY1 and indicated gRNA. α-tubulin was used as the loading control and each gRNA was normalized to effector-matched CIRTS expressed with NT gRNA for quantitation. n = 4 biological replicates. (**F**) Full western blots used in **Fig. S6E** quantitation. (**G**) Full western blots used in **Fig. 6E** quantitation. All bar graph values are shown as mean ± SEM with data points. Statistical analysis was performed using unpaired two-tailed Student's *t* test vs. CIRTS-4GT3/NT gRNA (**B**). Statistical analyses were performed using one-way ANOVA with *post hoc* Dunnett's multiple comparisons test vs. CIRTS-NY1/NT gRNA (**D**) and (**E**). $^*P<0.05$. No asterisk = not significant.

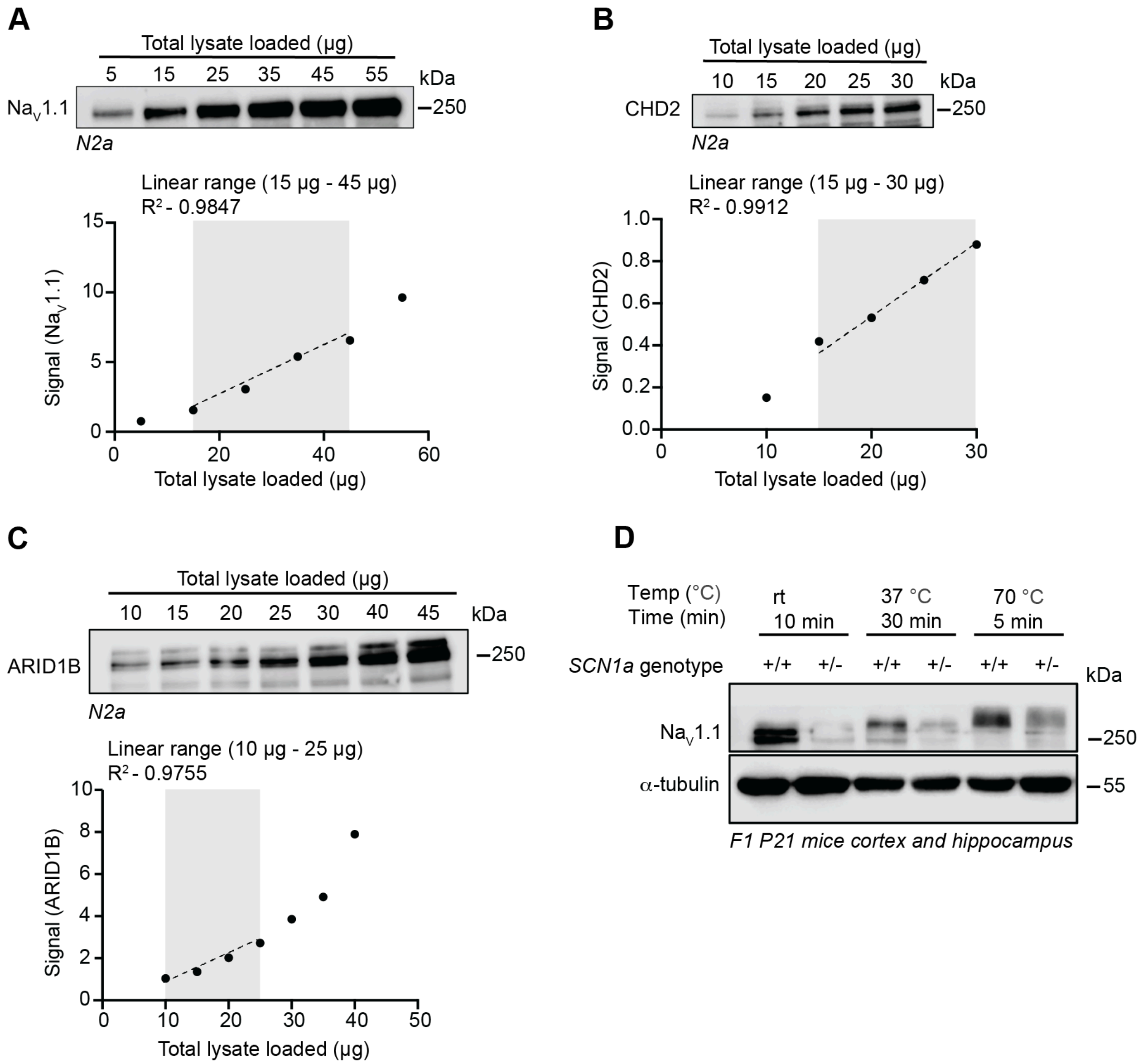


**Supplementary Figure 7**: Linear detection range for western blot primary antibodies and *in vivo* $Na_V1.1$ western blot optimization

(**A**) Western blot for $Na_V1.1$ over a range of total N2a lysate loaded. Signal for each band is plotted below with the total lysate loaded range with the highest linearity shaded. (**B**) Western blot for CHD2 over a range of total N2a lysate loaded. Signal for each band is plotted below with the total lysate loaded range with the highest linearity shaded. (**C**) Western blot for ARID1B over a range of total N2a lysate loaded. Signal for each band is plotted below with the total lysate loaded range with the highest linearity shaded. (**D**) Representative western blot for $SCN1a^{+/+}$ and $SCN1a^{+/-}$ F1 mouse cortex and hippocampus tissue samples blotted for $Na_V1.1$ and α-tubulin with different pre-loading sample heating steps. Statistical analyses were performed using simple linear regression to obtain $R^2$ for the dashed line fit to the indicated data points.

**Supplementary Table 1**: Representative plasmids used in this study

| # | Description | Stock ID | Name | Plasmid Map |
|---|---|---|---|---|
| 1 | CIRTS-NY1/NT | 58-48 | CIRTS-gRNA: cmv bdef-TBP6.7-NES-GGS-Y1 (1-364)-FLAG; hU6-1 TAR 1 stable NT gRNA | https://benchling.com/s/seq-QG6EmtgibQUUsqFz7ku8?m=slm-WKNA3fmuP5MiM3ppFJT2 |
| 2 | CIRTS-NY1/SCN1a-g1 | 72-129 | CIRTS-gRNA: cmv bdef-TBP6.7-NES-GGS-NY1-FLAG; hU6-1 TAR 1 stable mSCN1a g1 | https://benchling.com/s/seq-eh3xEwg05PcKmwvbUgkM?m=slm-EcA5r6n0AxXhvOftavAE |
| 3 | CIRTS-NY1/SCN1a-g2 | 72-130 | CIRTS-gRNA: cmv bdef-TBP6.7-NES-GGS-NY1-FLAG; hU6-1 TAR 1 stable mSCN1a g2 | https://benchling.com/s/seq-eHvRLz83IM9otTKKAeJS?m=slm-jKqUPQvjwYYuI7OoUmxX |
| 4 | CIRTS-NY1/SCN1a-g3 | 72-132 | CIRTS-gRNA: cmv bdef-TBP6.7-NES-GGS-NY1-FLAG; hU6-1 TAR 1 stable mSCN1a g3 | https://benchling.com/s/seq-SWjKDfHU9sBMzKz4RBSF?m=slm-j1oc5qpsaXsK46W5zx7h |
| 5 | CIRTS-NY1/SCN1a-g4 | 72-133 | CIRTS-gRNA: cmv bdef-TBP6.7-NES-GGS-NY1-FLAG; hU6-1 TAR 1 stable mSCN1a g4 | https://benchling.com/s/seq-TwBhongJ1acQx4BeEtBI?m=slm-i7B4fCkm7eP5DT3EfunC |
| 6 | CIRTS-NY1/SCN1a-g5 | 70-157 | CIRTS-gRNA: cmv bdef-TBP6.7-NES-GGS-NY1-FLAG; hU6-1 TAR 1 stable mSCN1a g5 | https://benchling.com/s/seq-esYT0lK4uy3XefMK4kU4?m=slm-ibeUIWeH65tGSCCUcNvH |
| 7 | CIRTS-4GT1/NT | 67-66 | CIRTS-gRNA: cmv bdef-TBP6.7-NES-GGS-eIF4GT1 (711-1599)-FLAG; hU6-1 TAR 1 stable NT gRNA | https://benchling.com/s/seq-QpRF9QT1QnWxYVxKLWaR?m=slm-6i8kKQNgQDQryQ4zcSOK |
| 8 | CIRTS-HSPB1/NT | 59-62 | CIRTS-gRNA: cmv bdef-TBP6.7-NES-GGS-HSPB1-FLAG; hU6-1 TAR 1 stable NT gRNA | https://benchling.com/s/seq-KfV0TE2blt1t8i8gehnB?m=slm-IJUbQ8Gm0KGGxYypiZot |
| 9 | CIRTS-eIF4e/NT | 42-42 | CIRTS-gRNA: cmv bdef-TBP6.7-NES-GGS-eIF4e-FLAG; hU6-1 TAR 1 stable NT gRNA | https://benchling.com/s/seq-x80ERqMg3D6PbUhsLr1c?m=slm-gY7obKaqnWWrxamr0CiL |

| | | | | |
|---|---|---|---|---|
| 10 | CIRTS-BolI/NT | 58-39 | CIRTS-gRNA: cmv bdef-TBP6.7-NES-GGS-BolI-FLAG; hU6-1 TAR 1 stable NT gRNA | https://benchling.com/s/seq-JU59ZWij2IYS9D8tL0cN?m=slm-2NNpVbpE7uS5yLUCdxJ6 |
| 11 | CIRTS-YB1/NT | 59-70 | CIRTS-gRNA: cmv bdef-TBP6.7-NES-GGS-YB1-FLAG; hU6-1 TAR 1 stable NT gRNA | https://benchling.com/s/seq-NpeQfdw8D4g9dLBstt5N?m=slm-Rp6Qd7vOsrrwYSXcLskd |
| 12 | CIRTS-HuR/NT | 70-39 | CIRTS-gRNA: cmv bdef-TBP6.7-NES-GGS-HuR-FLAG; hU6-1 TAR 1 stable NT gRNA | https://benchling.com/s/seq-w8b0naDt4e61d20eGULv?m=slm-35UHPBoAfEd0HznIpSIW |
| 13 | CIRTS-PCBP/NT | 70-41 | CIRTS-gRNA: cmv bdef-TBP6.7-NES-GGS-PCPB2-FLAG; hU6-1 TAR 1 stable NT gRNA | https://benchling.com/s/seq-3XQWQRCNuS0h6fkAoUwC?m=slm-Vqq33gFXCb9bQ7qCQJYG |
| 14 | CIRTS-eIF4a/NT | 58-43 | CIRTS-gRNA: cmv bdef-TBP6.7-NES-GGS-eIF4a-FLAG; hU6-1 TAR 1 stable NT gRNA | https://benchling.com/s/seq-TvjTkopM7ctHlgdQzME8?m=slm-lIYRBbqcdeQPmcqEIOmF |
| 15 | CIRTS-DAZ4/NT | 58-45 | CIRTS-gRNA: cmv bdef-TBP6.7-NES-GGS-DAZ4-FLAG; hU6-1 TAR 1 stable NT gRNA | https://benchling.com/s/seq-IFJIBnTQY3ZSNcUa3EI7?m=slm-j8mje99PDKcNNW4aM0be |
| 16 | CIRTS-FXR1/NT | 70-136 | CIRTS-gRNA: cmv bdef-TBP6.7-NES-GGS-FXR1-FLAG; hU6-1 TAR 1 stable NT gRNA | https://benchling.com/s/seq-XY7CXt0acrkUquIOBiGp?m=slm-NYRTEFIsakLGGRXeFLkA |
| 17 | CIRTS-PABPC1/NT | 58-44 | CIRTS-gRNA: cmv bdef-TBP6.7-NES-GGS-PABPC1-FLAG; hU6-1 TAR 1 stable NT gRNA | https://benchling.com/s/seq-TbbfZLODIeGelwYLNDFY?m=slm-NqoUSsrPqYwhZVp14T9X |
| 18 | CIRTS-4GT1/SCN1a-g6 | 73-44 | CIRTS-gRNA: cmv bdef-TBP6.7-NES-GGS-eIF4GT1 (711-1599)-FLAG; hU6-1 TAR 1 stable g6 5' UTR mSCN1a gRNA | https://benchling.com/s/seq-J5qWo5GyCYwRi1IokToB?m=slm-SnwUJ1OKM1BSsBGidgfy |
| 19 | CIRTS-4GT1/SCN1a-g7 | 73-47 | CIRTS-gRNA: cmv bdef-TBP6.7-NES-GGS-eIF4GT1 (711-1599)-FLAG; hU6-1 TAR 1 stable g7 5' UTR mSCN1a gRNA | https://benchling.com/s/seq-GStpypNzJk8DHn0AICDd?m=slm-Ip5ppEA5SBngUGjqzDvb |

| | | | | |
|---|---|---|---|---|
| 20 | CIRTS-GFP/SCN1a-g1 | 76-139 | CIRTS-gRNA: cmv bdef-TBP6.7-NES-GGS-GFP-FLAG; hU6-1 TAR 1 stable g1 3' UTR mSCN1a gRNA | https://benchling.com/s/seq-2J9kTrr8ga6ki2pYWzMi?m=slm-0dzdQvM4ptvLiZiVBQGt |
| 21 | CIRTS-4GT2/SCN1a-g1 | 76-121 | 76-121: CIRTS-gRNA: cmv bdef-TBP6.7-NES-GGS-eIF4GT2 (539-1105)-FLAG; hU6-1 TAR 1 stable g1 3' UTR mSCN1a gRNA | https://benchling.com/s/seq-ifgnhiKhWyL9c2iaNg5R?m=slm-IeLMhzEJLY7c6dwkpYHH |
| 22 | CIRTS-4GT3/SCN1a-g1 | 72-166 | CIRTS-gRNA: cmv bdef-TBP6.7-NES-GGS-eIF4GT3 (711-1105)-FLAG; hU6-1 TAR 1 stable g1 3' UTR mSCN1a gRNA | https://benchling.com/s/seq-xn2fvGy2rj7h4wHOji3P?m=slm-pScOO4ueLJqgSmusLcPR |
| 23 | CIRTS-4GT4/SCN1a-g1 | 76-120 | CIRTS-gRNA: cmv bdef-TBP6.7-NES-GGS-eIF4GT4 (1011-1105)-FLAG; hU6-1 TAR 1 stable g1 3' UTR mSCN1a gRNA | https://benchling.com/s/seq-nnnBwPJJgktMDksjJYUM?m=slm-Bx90P8m3wUgZRlPu3j3E |
| 24 | Empty CIRTS-gRNA Vector | 70-44 | Empty CIRTS-gRNA: cmv_bGH polyA; hU6-TTTTTTT | https://benchling.com/s/seq-AwuTv3oP1ffGuqQ0JDAm?m=slm-NxVI3zJr7pjd8C83yI3T |
| 25 | CIRTS-NY1/CHD2-g1 | 76-192 | CIRTS-gRNA: cmv bdef-TBP6.7-NES-GGS-eIF4GT3 (711-1105)-FLAG; hU6-1 TAR 1 stable g1 mCHD2 gRNA | https://benchling.com/s/seq-oodmxf1PmeIH51MMwi8S?m=slm-GcWKfgBo3w5G3B3ygnfP |
| 26 | CIRTS-NY1/CHD2-g2 | 76-193 | CIRTS-gRNA: cmv bdef-TBP6.7-NES-GGS-eIF4GT3 (711-1105)-FLAG; hU6-1 TAR 1 stable g2 mCHD2 gRNA | https://benchling.com/s/seq-UXeaf3qoGbyDsYXEm5gK?m=slm-59TkZoRgBqhncCmC2gpR |
| 27 | CIRTS-NY1/CHD2-g3 | 76-194 | CIRTS-gRNA: cmv bdef-TBP6.7-NES-GGS-eIF4GT3 (711-1105)-FLAG; hU6-1 TAR 1 stable g3 mCHD2 gRNA | https://benchling.com/s/seq-H4hu6fifNXFr4DtzTqWo?m=slm-MC3cLMUWaAZbIVdFW4Eb |
| 28 | CIRTS-NY1/CHD2-g4 | 76-195 | CIRTS-gRNA: cmv bdef-TBP6.7-NES-GGS-eIF4GT3 (711-1105)-FLAG; hU6-1 TAR 1 stable g4 mCHD2 gRNA | https://benchling.com/s/seq-IJX7K8N9MDSdrQZLIMLn?m=slm-bye7j5KomTREWZ6spuvU |
| 29 | CIRTS-NY1/CHD2-g5 | 76-196 | CIRTS-gRNA: cmv bdef-TBP6.7-NES-GGS-eIF4GT3 (711-1105)-FLAG; hU6-1 TAR 1 stable g5 mCHD2 gRNA | https://benchling.com/s/seq-8L9qhqg8DDZF15ZEubkk?m=slm-l9lSaVfG3InA1AAO7x9I |

| | | | | |
|---|---|---|---|---|
| 30 | Lentiviral vector: CIRTS-4GT3/NT | 76-66 | lenti-hU6::hU6-NT gRNA 1 TAR 1 Stable; EF1a-GCCACC-Bdef-TBP-eIF4GT3 (711-1105)-P2A-Puro | https://benchling.com/s/seq-xMx2yIOwvY1zfnAYWWGx?m=slm-0O00dXS1He4t3TZ3ZWhk |
| 31 | Lentiviral vector: CIRTS-4GT3/SCN1a-g1 | 76-67 | lenti-hU6::hU6-g1 SCN1a gRNA 1 TAR 1 Stable; EF1a-GCCACC-Bdef-TBP-eIF4GT3 (711-1105)-P2A-Puro | https://benchling.com/s/seq-3Gp5T7ZI59l4pFIFoO5d?m=slm-kWKxA168tPDnAPJAnsuZ |
| 32 | Lentiviral vector: CIRTS-NY1/NT | 72-172 | 72-172: lenti-hU6::hU6-NT gRNA 1 TAR 1 Stable; EF1a-GCCACC-Bdef-TBP-Y1 (1-364)-P2A-Puro | https://benchling.com/s/seq-GkOLrVti7GbxZOiCj0FM?m=slm-UZcdszzO7Bo9So9iL8aT |
| 33 | Lentiviral vector: CIRTS-NY1/rSCN1a-g5 | 74-36 | 74-36: lenti-hU6::hU6-rat SCN1a g5 gRNA 1 TAR 1 Stable; EF1a-GCCACC-Bdef-TBP-Y1 (1-364)-P2A-Puro | https://benchling.com/s/seq-yE5I5Prr36kBD9kc74uS?m=slm-yDTBxMfnFgnYEFGSojUW |
| 34 | AAV vector: CIRTS-4GT3/SCN1a-g1 | 77-171 | pX601-AAV-hSYN::bdef-TBP6.7-NES-GGS-eIF4GT3 (711-1105)-FLAG; hU6-1 TAR 1 stable g1 mSCN1a gRNA | https://benchling.com/s/seq-75KIuFkPYdc3qbJmOq16?m=slm-7T6kuQGWmlt5tCNZKydC |
| 35 | Dual luciferase reporter with mSCN1a 3’ UTR | 71-24 | pmirGLO-PGK-CACC-Fluc-mSCN1a 3' UTR SV40-NLuc (CP1-PEST) | https://benchling.com/s/seq-AyKP85keBVWg5S5smCk9?m=slm-XvrTiRMaiHMC6JVPffeb |
| 36 | Dual luciferase reporter with mCHD2 3’ UTR | 74-39 | pmirGLO-PGK-CACC-Fluc-mCHD2 3' UTR SV40-NLuc (CP1-PEST) | https://benchling.com/s/seq-v8qtqGJ80SiwnJwbPrFP?m=slm-rgX3NWJo2eRww4S75IQ3 |

**Supplementary Table 2:** CIRTS gRNA sequences

| # | Target-Number: | Sequence (5' to 3'): |
|---|---|---|
| 1 | SCN1a-g1 | GACUUCACAGGCUGUAAACAAUUUGUCACCCAAUUUAUUA |
| 2 | SCN1a-g2 | GUCACAGAGUUUGCCGACAAGGGGGUCACUGUCUUAUUGA |
| 3 | SCN1a-g3 | AUUUAUAAAGGGUUCUUUUGGUCAAUUCGCUCUGCUAGGG |
| 4 | SCN1a-g4 | AUUUCUUAGUUAGCAAAUAUCUUCCAUGAAAAACUGCUGU |
| 5 | SCN1a-g5 | CUCUCUGAUUUUGAGGAAAAGAAAUUCUGAAGCCCACAUU |
| 6 | SCN1a-g6 | AACAGUUCCAUGAGUUUCCACAGGCCGGUAGUGAGUCUCC |
| 7 | SCN1a-g7 | CAUGUGAGAUUCCCCCGAAAGAUGUAUGUAGUUAAAGCUC |
| 8 | CHD2-g1 | GAUUCAACAGCAGCAGCAUAUCCAGAAACCCGGCUCAGCA |
| 9 | CHD2-g2 | UGUAUCAUUUCUCCCAAUAACACAGCUAAGGGCUAAGGGG |
| 10 | CHD2-g3 | CCAACCCAACCCAACCCAACCCGACGCUGACCAGCCUCCU |
| 11 | CHD2-g4 | GCUUCCUUUCUUUCAACCCCUCUUCAAAUCCACUCCCUCA |
| 12 | CHD2-g5 | CAAGUGGGUUUGUUUGGUCUGUAGGAUUUCAGUGUUGCUG |
| 13 | SCN1a-rg5 | GAGGAAAAGAAAUUCCGAAGCUCACGUUUAUAUUUAGAAC |
| 14 | ARID1B-g1 | UAAUUCCCAAUUGGUAAAUAGUAAACGGGGCAGAGGAGCA |
| 15 | ARID1B-g2 | AAAAAAAGAAACAGAAAACCGCCAGAGGAUGUACUUAUAC |
| 16 | ARID1B-g3 | GAACUGAGCCUGUCUAUACCUUUCAUUCAUUGUCUGCAAG |

**Supplementary Table 3:** CIRTS effectors amino acid sequences

| # | Name | Amino acid sequence |
|---|---|---|
| 1 | NY1 (1-364) | MSATSVDTQRTKGQDNKVQNGSLHQKDTVHDNDFEPYLTGQ<br>SNQSNSYPSMSDPYLSSYYPPSIGFPYSLNEAPWSTAGDPPI<br>PYLTTYGQLSNGDHHFMHDAVFGQPGGLGNNIYQHRFNFFP<br>ENPAFSAWGTSGSQGQQTQSSAYGSSYTYPPSSLGGTVVDG<br>QPGFHSDTLSKAPGMNSLEQGMVGLKIGDVSSSAVKTVGSVV<br>SSVALTGVLSGNGGTNVNMPVSKPTSWAAIASKPAKPQPKMK<br>TKSGPVMGGGLPPPPIKHNMDIGTWDNKGPVPKAPVPQQAP<br>SPQAAPQPQQVAQPLPAQPPALAQPQYQSPQQPPQTRWVAP<br>RNRNAAFGQSGGAGSDSNSPGNVQPNSAPSVES |
| 2 | eIF4GI-T1<br>(711-1599) | EPRKIIATVLMTEDIKLNKAEKAWKPSSKRTAADKDRGEEDAD<br>GSKTQDLFRRVRSILNKLTPQMFQQLMKQVTQLAIDTEERLKG<br>VIDLIFEKAISEPNFSVAYANMCRCLMALKVPTTEKPTVTVNFR<br>KLLLNRCQKEFEKDKDDDEVFEKKQKEMDEAATAEERGRLKE<br>ELEEARDIARRRSLGNIKFIGELFKLKMLTEAIMHDCVVKLLKN<br>HDEESLECLCRLLTTIGKDLDFEKAKPRMDQYFNQMEKIIKEK<br>KTSSRIRFMLQDVLDLRGSNWVPRRGDQGPKTIDQIHKEAEM<br>EEHREHIKVQQLMAKGSDKRRGGPPGPPISRGLPLVDDGGW<br>NTVPISKGSRPIDTSRLTKITKPGSIDSNNQLFAPGGRLSWGK<br>GSSGGSGAKPSDAASEAARPATSTLNRFSALQQAVPTESTDN<br>RRVVQRSSLSRERGEKAGDRGDRLERSERGGDRGDRLDRA<br>RTPATKRSFSKEVEERSRERPSQPEGLRKAASLTEDRDRGRD<br>AVKREAALPPVSPLKAALSEEELEKKSKAIIEEYLHLNDMKEAV<br>QCVQELASPSLLFIFVRHGVESTLERSAIAREHMGQLLHQLLC<br>AGHLSTAQYYQGLYEILELAEDMEIDIPHVWLYLAELVTPILQEG<br>GVPMGELFREITKPLRPLGKAASLLLEILGLLCKSMGPKKVGTL<br>WREAGLSWKEFLPEGQDIGAFVAEQKVEYTLGEESEAPGQR<br>ALPSEELNRQLEKLLKEGSSNQRVFDWIEANLSEQQIVSNTLV<br>RALMTAVCYSAIIFETPLRVDVAVLKARAKLLQKYLCDEQKELQ<br>ALYALQALVVTLEQPPNLLRMFFDALYDEDVVKEDAFYSWESS<br>KDPAEQQGKGVALKSVTAFFKWLREAEEESDHN |
| 3 | eIF4aI | MSASQDSRSRDNGPDGMEPEGVIESNWNEIVDSFDDMNLSE<br>SLLRGIYAYGFEKPSAIQQRAILPCIKGYDVIAQAQSGTGKTATF<br>AISILQQIELDLKATQALVLAPTRELAQQIQKVVMALGDYMGAS<br>CHACIGGTNVRAEVQKLQMEAPHIIVGTPGRVFDMLNRRYLS<br>PKYIKMFVLDEADEMLSRGFKDQIYDIFQKLNSNTQVVLLSAT<br>MPSDVLEVTKKFMRDPIRILVKKEELTLEGIRQFYINVEREEWK<br>LDTLCDLYETLTITQAVIFINTRRKVDWLTEKMHARDFTVSAMH<br>GDMDQKERDVIMREFRSGSSRVLITTDLLARGIDVQQVSLVIN<br>YDLPTNRENYIHRIGRGGRFGRKGVAINMVTEEDKRTLRDIET<br>FYNTSIEEMPLNVADLI |
| 4 | PABPC1 | MNPSAPSYPMASLYVGDLHPDVTEAMLYEKFSPAGPILSIRVC<br>RDMITRRSLGYAYVNFQQPADAERALDTMNFDVIKGKPVRIM<br>WSQRDPSLRKSGVGNIFIKNLDKSIDNKALYDTFSAFGNILSCK |

| | | |
|---|---|---|
| | | VVCDENGSKGYGFVHFETQEAAERAIEKMNGMLLNDRKVFV<br>GRFKSRKEREAELGARAKEFTNVYIKNFGEDMDDERLKDLFG<br>KFGPALSVKVMTDESGKSKGFGFVSFERHEDAQKAVDEMNG<br>KELNGKQIYVGRAQKKVERQTELKRKFEQMKQDRITRYQGVN<br>LYVKNLDDGIDDERLRKEFSPFGTITSAKVMMEGGRSKGFGF<br>VCFSSPEEATKAVTEMNGRIVATKPLYVALAQRKEERQAHLTN<br>QYMQRMASVRAVPNPVINPYQPAPPSGYFMAAIPQTQNRAAY<br>YPPSQIAQLRPSPRWTAQGARPHPFQNMPGAIRPAAPRPPFS<br>TMRPASSQVPRVMSTQRVANTSTQTMGPRPAAAAAAATPAVR<br>TVPQYKYAAGVRNPQQHLNAQPQVTMQQPAVHVQGQEPLTA<br>SMLASAPPQEQKQMLGERLFPLIQAMHPTLAGKITGMLLEIDN<br>SELLHMLESPESLRSKVDEAVAVLQAHQAKEAAQKAVNSATG<br>VPTV |
| 5 | eIF4e | MATVEPETTPTPNPPTTEEEKTESNQEVANPEHYIKHPLQNR<br>WALWFFKNDKSKTWQANLRLISKFDTVEDFWALYNHIQLSSNL<br>MPGCDYSLFKDGIEPMWEDEKNKRGGRWLITLNKQQRRSDL<br>DRFWLETLLCLIGESFDDYSDDVCGAVVNVRAKGDKIAIWTTE<br>CENREAVTHIGRVYKERLGLPPKIVIGYQSHADTATKSGSTTKN<br>RFVV |
| 6 | YB-1 | MSSEAETQQPPAAPPAAPALSAADTKPGTTGSGAGSGGPGG<br>LTSAAPAGGDKKVIATKVLGTVKWFNVRNGYGFINRNDTKEDV<br>FVHQTAIKKNNPRKYLRSVGDGETVEFDVVEGEKGAEAANVT<br>GPGGVPVQGSKYAADRNHYRRYPRRRGPPRNYQQNYQNSE<br>SGEKNEGSESAPEGQAQQRRPYRRRRFPPYYMRRPYGRRP<br>QYSNPPVQGEVMEGADNQGAGEQGRPVRQNMYRGYRPRF<br>RRGPPRQRQPREDGNEEDKENQGDETQGQQPPQRRYRRN<br>FNYRRRRPENPKPQDGKETKAADPPAENSSAPEAEQGGAE |
| 7 | HSPB1 | MTERRVPFSLLRGPSWDPFRDWYPHSRLFDQAFGLPRLPEE<br>WSQWLGGSSWPGYVRPLPPAAIESPAVAAPAYSRALSRQLSS<br>GVSEIRHTADRWRVSLDVNHFAPDELTVKTKDGVVEITGKHEE<br>RQDEHGYISRCFTRKYTLPPGVDPTQVSSSLSPEGTLTVEAP<br>MPKLATQSNEITIPVTFESRAQLGGPEAAKSDETAAKL |
| 8 | Boll | MQTDSLSPSPNPVSPVPLNNPTSAPRYGTVIPNRIFVGGIDFK<br>TNESDLRKFFSQYGSVKEVKIVNDRAGVSKGYGFVTFETQED<br>AQKILQEAEKLNYKDKKLNIGPAIRKQQVGIPRSSIMPAAGTMY<br>LTTSTGYPYTYHNGVAYFHTPEVTSVPPPWPSRSVCSSPVMV<br>AQPIYQQPAYHYQATTQYLPGQWQWSVPQPSASSAPFLYLQ<br>PSEVIYQPVEIAQDGGCVPPPLSLMETSVPEPYSDHGVQATY<br>HQVYAPSAITMPAPVMQPEPIKTVWSIHY |
| 9 | PCBP2 | MDTGVIEGGLNVTLTIRLLMHGKEVGSIIGKKGESVKKMREES<br>GARINISEGNCPERIITLAGPTNAIFKAFAMIIDKLEEDISSSMTN<br>STAASRPPVTLRLVVPASQCGSLIGKGGCKIKEIRESTGAQVQ<br>VAGDMLPNSTERAITIAGIPQSIIECVKQICVVMLESPPKGVTIP<br>YRPKPSSSPVIFAGGQAYTIQGQYAIPQPDLTKLHQLAMQQSH<br>FPMTHGNTGFSGIESSSPEVKGYWAGLDASAQTTSHELTIPN |

| | | |
|---|---|---|
| | | DLIGCIIGRQGAKINEIRQMSGAQIKIANPVEGSTDRQVTITGSA ASISLAQYLINVRLSSETGGMGSS |
| 10 | HuR (ELAVL1) | MSNGYEDHMAEDCRGDIGRTNLIVNYLPQNMTQDELRSLFSS IGEVESAKLIRDKVAGHSLGYGFVNYVTAKDAERAINTLNGLRL QSKTIKVSYARPSSEVIKDANLYISGLPRTMTQKDVEDMFSRF GRIINSRVLVDQTTGLSRGVAFIRFDKRSEAEEAITSFNGHKPP GSSEPITVKFAANPNQNKNVALLSQLYHSPARRFGGPVHHQA QRFRFSPMGVDHMSGLSGVNVPGNASSGWCIFIYNLGQDAD EGILWQMFGPFGAVTNVKVIRDFNTNKCKGFGFVTMTNYEEA AMAIASLNGYRLGDKILQVSFKTNKSHK |
| 11 | FXR-1 | MAELTVEVRGSNGAFYKGFIKDVHEDSLTVVFENNWQPERQV PFNEVRLPPPPDIKKEISEGDEVEVYSRANDQEPCGWWLAKV RMMKGEFYVIEYAACDATYNEIVTFERLRPVNQNKTVKKNTFF KCTVDVPEDLREACANENAHKDFKKAVGACRIFYHPETTQLMI LSASEATVKRVNILSDMHLRSIRTKLMLMSRNEEATKHLECTK QLAAAFHEEFVVREDLMGLAIGTHGSNIQQARKVPGVTAIELD EDTGTFRIYGESADAVKKARGFLEFVEDFIQVPRNLVGKVIGK NGKVIQEIVDKSGVVRVRIEGDNENKLPREDGMVPFVFVGTK ESIGNVQVLLEYHIAYLKEVEQLRMERLQIDEQLRQIGSRSYSG RGRGRRGPNYTSGYGTNSELSNPSETESERKDELSDWSLAG EDDRDSRHQRDSRRRPGGRGRSVSGGRGRGGPRGGKSSIS SVLKDPDSNPYSLLDNTESDQTADTDASESHHSTNRRRRSRR RRTDEDAVLMDGMTESDTASVNENGLGKRCD |
| 12 | DAZ4 | MSAANPETPNSTISREASTQSSSAAASQGWVLPEGKIVPNTV FVGGIDARMDETEIGSCFGRYGSVKEVKIITNRTGVSKGYGFV SFVNDVDVQKIVGSQIHFHGKKLKLGPAIRKQKLCARHVQPRP LVVNPPPPPQFQNVWRNPNTETYLQPQITPNPVTQHVQAYSA YPHSPGQVITGCQLLVYNYQEYPTYPDSAFQVTTGYQLPVYN YQPFPAYPRSPFQVTAGYQLPVYNYQAFPAYPNSPFQVATGY QFPVYNYQPFPAYPSSPFQVTAGYQLPVYNYQAFPAYPNSPF QVATGYQFPVYNYQAFPAYPNSPVQVTTGYQLPVYNYQAFPA YPSSPFQVTTGYQLPVYNYQAFPAYPNSAVQVTTGYQFHVYN YQMPPQCPVGEQRRNLWTEAYKWWYLVCLIQRRD |
| 13 | eIF4GI-T2 (539-1104) | PAVPEVENQPPAGSNPGPESEGSGVPPRPEEADETWDSKED KIHNAENIQPGEQKYEYKSDQWKPLNLEEKKRYDREFLLGFQ FIFASMQKPEGLPHISDVVLDKANKTPLRPLDPTRLQGINCGP DFTPSFANLGRTTLSTRGPPRGGPGGELPRGPAGLGPRRSQ QGPRKEPRKIIATVLMTEDIKLNKAEKAWKPSSKRTAADKDRG EEDADGSKTQDLFRRVRSILNKLTPQMFQQLMKQVTQLAIDTE ERLKGVIDLIFEKAISEPNFSVAYANMCRCLMALKVPTTEKPTV TVNFRKLLLNRCQKEFEKDKDDDEVFEKKQKEMDEAATAEER GRLKEELEEARDIARRRSLGNIKFIGELFKLKMLTEAIMHDCVV KLLKNHDEESLECLCRLLTTIGKDLDFEKAKPRMDQYFNQMEK IIKEKKTSSRIRFMLQDVLDLRGSNWVPRRGDQGPKTIDQIHK EAEMEEHREHIKVQQLMAKGSDKRRGGPPGPPISRGLPLVDD GGWNTVPISKGSRPIDTSRLTKITKPGSIDSNNQLFAPGGRLS |

| | | |
|---|---|---|
| | | WGKGSSGGSGAK |
| 14 | eIF4GI-T3 (711-1104) | EPRKIIATVLMTEDIKLNKAEKAWKPSSKRTAADKDRGEEDAD GSKTQDLFRRVRSILNKLTPQMFQQLMKQVTQLAIDTEERLKG VIDLIFEKAISEPNFSVAYANMCRCLMALKVPTTEKPTVTVNFR KLLLNRCQKEFEKDKDDDEVFEKKQKEMDEAATAEERGRLKE ELEEARDIARRRSLGNIKFIGELFKLKMLTEAIMHDCVVKLLKN HDEESLECLCRLLTTIGKDLDFEKAKPRMDQYFNQMEKIIKEK KTSSRIRFMLQDVLDLRGSNWVPRRGDQGPKTIDQIHKEAEM EEHREHIKVQQLMAKGSDKRRGGPPGPPISRGLPLVDDGGW NTVPISKGSRPIDTSRLTKITKPGSIDSNNQLFAPGGRLSWGK GSSGGSGAK |
| 15 | eIF4GI-T4 (1011-1104) | MEEHREHIKVQQLMAKGSDKRRGGPPGPPISRGLPLVDDGG WNTVPISKGSRPIDTSRLTKITKPGSIDSNNQLFAPGGRLSWG KGSSGGSGAK |

**Supplementary Table 4:** Primer sequences

| # | Target (Assay) | Sequence (5' to 3'): |
|---|---|---|
| 1 | mSCN1a-fwd (qPCR) | GGTCATGGTGATTGGGAACCTTG |
| 2 | mSCN1a-rev (qPCR) | CATCCTGTCCACAGCAATCTGC |
| 3 | mGAPDH-fwd (qPCR) | CATCACTGCCACCCAGAAGACTG |
| 4 | mGAPDH-rev (qPCR) | ATGCCAGTGAGCTTCCCGTTCAG |
| 5 | mCHD2-fwd (qPCR) | GGAGATCATAGAACGGGCCA |
| 6 | mCHD2-rev (qPCR) | AAAAGGGTTTGAGTTGGATCTTC |
| 7 | mSCN1a common fwd (Genotyping) | AGTCTGTACCAGGCAGAACTTG |
| 8 | mSCN1a wild type rev (Genotyping) | CCCTGAGATGTGGGTGAATAG |
| 9 | mSCN1a knockout rev (Genotyping) | AGACTGCCTTGGGAAAAGCG |

**Supplementary Table 5:** Antibodies and dilutions used for western blots

| # | Target: | Vendor | Catalogue number | Dilution |
|---|---|---|---|---|
| 1 | SCN1a ($Na_V$1.1) | Alomone lab | ASC-001 | 1:500 |
| 2 | α-tubulin (HRP) | Proteintech | HRP-66031 | 1:5000 |
| 3 | CHD2 | Cell Signaling Tech | CST-4170 | 1:1000 |
| 4 | ARID1B | Cell Signaling Tech | CST-92964 | 1:1000 |
| 5 | DYKDDDDK (FLAG tag) | Invitrogen | MA1-91878 | 1:500 |
| 6 | mouse IgG H&L (HRP) | Abcam | ab6721 | 1:5000 |
| 7 | rabbit IgG H&L (HRP) | Abcam | ab6728 | 1:5000 |
